# Systems-level analysis of RDK1 reveals compartment-specific kinase activity and a function in the maintenance of the mitochondrial proteome in *Trypanosoma brucei*

**DOI:** 10.64898/2026.05.05.722970

**Authors:** Ashutosh P. Dubey, Parul Pandey, Dinh Song H. Bui, Chukwujekwu O. Aleke, Joseph T. Smith

**Affiliations:** Department of Microbiology and Immunology, Jacobs School of Medicine and Biomedical Sciences, University at Buffalo, Buffalo, New York, USA

**Keywords:** Trypanosoma brucei, kinase, mitochondria, signaling, differentiation

## Abstract

Repressor-of-differentiation kinase 1 (RDK1) is one of two kinases expressed in bloodstream form *Trypanosoma brucei* parasites that were found to repress premature and spontaneous differentiation into the insect procyclic form. However, the effect of RDK1 RNAi was previously limited to the expression of a single surface coat protein, EP1 procyclin. Thus, there remains a significant gap in knowledge on the impact of RDK1 expression in bloodstream form *T. brucei* parasites. Here, we employ a systems biology approach and performed several proteomics analyses to identify RDK1 protein interactions and to determine the impact of loss of RDK1 expression on the bloodstream form proteome and phosphoproteome to uncover clues about potential mechanisms for RDK1 function. We found that RDK1 is dual localized to the cell membrane and the mitochondrial inner membrane with the kinase domain oriented towards the cytoplasm and mitochondrial inner membrane. Unexpectedly, the most enriched RDK1-proximal proteins were mitochondrial proteins. Furthermore, RDK1 depletion causes bloodstream form parasites to significantly upregulate many mitochondrial proteins and glycosomal proteins, several of which are upregulated in procyclic form parasites. Surprisingly, the mitochondrial phosphoproteome is largely unaffected by RDK1 depletion, while RDK1-dependent phosphoregulation is restricted to the cell membrane localization of RDK1. Lastly, we determined that RDK1 does not possess adenyl cyclase activity or alter intracellular cAMP levels; however, the dysregulated phosphoproteins correlate with functions in cyclic nucleotide signaling. In conclusion, RDK1 exhibits localization-specific kinase activity to regulate cyclic nucleotide signaling and mitochondrial proteomic maintenance in bloodstream form parasites.

**IMPORTANCE:** *Trypanosoma brucei* is the unicellular parasite that causes African sleeping sickness and nagana disease in livestock across 36 sub-Saharan African countries. The parasite encounters different environmental niches as it is transmitted from an infected human to the tsetse fly vector as the fly takes a blood meal. *T. brucei* must sense environmental cues to initiate intracellular signaling pathways to promote effective differentiation and cellular remodeling from the mammalian bloodstream forms to the insect procyclic form. RDK1 is one of two kinases shown to repress premature differentiation to procyclic form, which would be detrimental for parasite survival in the human host. Therefore, it is essential to uncover mechanisms of RDK1 function to better understand how *T. brucei* maintains homeostasis in the human host and signals for effective cellular remodeling during parasite transmission.

## INTRODUCTION

*Trypanosoma brucei* is the protozoan parasite that causes African sleeping sickness in humans and nagana disease in domesticated livestock and pets across sub-Saharan Africa (1, 2). *T. brucei* is transmitted by the bite of infected tsetse flies as they take a blood meal from a mammalian host (3). In the mammalian host, *T. brucei* multiplies as the slender bloodstream form (BSF) and maintains infection of multiple host tissues such as the blood, skin, adipose tissue, lymph, and cerebrospinal fluid (4–7). When parasitemia in host tissue niches are high, the subpopulation of slender BSF parasites irreversibly differentiate into the quiescent stumpy BSF, which are molecularly pre-adapted for differentiation to the procyclic form (PCF) in the midgut of the tsetse fly following a blood meal (3, 8). This transmission event coincides with various environmental shifts such as a notable temperature reduction from 37°C to 27°C, glucose availability, pH, and concentrations of various metabolites (9–12). Environmental changes can serve as differentiation cues that trigger intracellular signaling pathways that promote effective parasite differentiation (3, 13–16).

Protein phosphorylation is one of the most studied eukaryotic post-translational protein modifications. In *T. brucei*, protein phosphorylation has been shown to be involved in parasite environmental stress responses, cell cycle regulation, and life cycle progression (13, 17–21). Yet there is little mechanistic insight on the kinases involved in maintaining inhibitory phosphorylation events in the BSF proteome that work to inhibit PCF differentiation. TbMAPK5, TbDYRK, and YAK are three kinases that are involved in the regulation of slender-to-stumpy BSF differentiation. While genetic knockout (KO) of TbMAPK5 causes accelerated slender-to-stumpy BSF differentiation at lower parasite densities (22), TbDYRK KO or YAK KO in slender BSF parasites completely prevents stumpy BSF formation (16, 23). In contrast, depletion by RNA interference (RNAi) of NRKA and NRKB in slender BSF parasites does not impact stumpy BSF formation; however, NRKA/NRKB RNAi strongly inhibits BSF-to-PCF differentiation upon *in vitro* differentiation stimulation (19). However, it is notable that genetic KO or RNAi of these kinases does not initiate BSF-to-PCF differentiation in the absence of *in vitro* differentiation stimulus. To date, only two kinases have been identified that, when depleted by RNAi, initiate BSF-to-PCF differentiation in the absence of *in vitro* differentiation cues and have been named <u>r</u>epressor-of-<u>d</u>ifferentiation <u>k</u>inase <u>1</u> and <u>2</u> (RDK1 and RDK2) (17). While RDK1 RNAi or RDK2 RNAi causes a subpopulation of the BSF parasites to acquire PCF-like morphology and express the PCF surface coat protein EP1 procyclin, only RDK1 RNAi parasites increase the percentage of parasites that express EP1 procyclin when exposed to various differentiation cues, such as a reduction in temperature from 37°C to 27°C, cAMP treatment, or treatment with the phosphatase inhibitor BZ3 (17). Thus, RDK1 function likely represses differentiation signaling pathways that are sensitive to differentiation cues.

RDK1 is a membrane-integrated STE-like kinase that has three predicted protein domains: an N-terminal ligand-binding (or bacterial sensor-like) domain, a central adenyl/guanyl cyclase domain, and a C-terminal STE-like kinase domain (17, 24). The domain organization of RDK1 resembles receptor guanyl cyclases in metazoans (25, 26). However, canonical receptor guanyl cyclases often lack *in vivo* catalytic kinase activity due to the lack of catalytic residues or ATP binding to the kinase homology domain induces conformational changes that alters protein substrate specificity or activity, possibly precluding the *in vivo* phosphorylation of substrates (27–29). In contrast, RDK1 has an inverted organization of the cyclase and kinase domains that may permit functional kinase activity. This raises the intriguing possibility that RDK1 integrates ligand sensing with dual enzymatic signaling in a noncanonical manner where kinase activity may be active *in vivo*.

To date, very little research on RDK1 function has been performed. Thus far, the only published phenotypes of BSF parasites that have been depleted of RDK1 are: RDK1 is implicated in assisting to regulate the cell cycle in BSF parasites (17), RDK1 RNAi parasites increase *PAD1* mRNA expression (17), EP1 procyclin is expressed in a subset of RDK1 RNAi parasites that is further enhanced by differentiation cues (17), procyclin-positive RDK1 RNAi parasites have a PCF-like morphology with a repositioned kinetoplast (17), the RDK1 RNAi transcriptome implicates the initiation of a differentiation program (17), and RDK1 RNAi induces the editing of mitochondrial-encoded *COII* mRNA that is further enhanced by a reduction in temperature (30). However, there is a substantial gap in knowledge on the impact of RDK1 on the BSF proteome and, more specifically, the BSF phosphoproteome. The primary goal of our study was to take a systems-level approach to determine RDK1 localization, protein interactions, and the impact of RDK1 depletion on the abundances of parasite proteins and phosphopeptides. Herein, we discovered that RDK1 dual-localizes both the cell membrane and the mitochondrial inner membrane. Furthermore, despite RDK1 having the most abundant interactions with many mitochondrial proteins and RDK1 depletion having a significant impact on the mitochondrial proteome, RDK1 does not impact the mitochondrial phosphoproteome. Instead, RDK1-dependent phosphorylation events are restricted primarily to its cell membrane localization, suggesting that RDK1 kinase activity is cell localization-specific.

## RESULTS

### RDK1 expression is downregulated and is not essential for cell growth in PCF parasites

Prior to this work, relative RDK1 protein expression between BSF and PCF parasites had not been tested directly, and previously published omics studies offer a conflicting conclusion. RNA-Seq and ribosome profiling studies suggest that *RDK1* mRNA abundance and ribosome occupancy are increased in BSF relative to PCF (31, 32). In contrast, a quantitative proteomics study that evaluated protein expression profile patterns as *T. brucei* progressively differentiates from BSF to PCF indicates that RDK1 is unexpectedly upregulated in PCF compared to BSF (33). This has led to the speculation that RDK1 continues to be expressed in PCF parasites to carry out an unknown function. Therefore, we wanted to clarify RDK1 expression between BSF and PCF in a pleomorphic *T. brucei* strain. To directly determine *RDK1* mRNA levels as BSF parasites differentiate into PCF, we isolated RNA from slender BSF, stumpy BSF, stumpy BSF exposed to PCF differentiation stimuli for 24 hours (hereafter referred to as “differentiating” parasites), and PCF parasites to perform quantitative RT-PCR analysis (Fig. 1A). We found that *RDK1* mRNA levels are increased 1.80-fold in stumpy BSF relative to slender BSF parasites. However, when stumpy BSF are stimulated to differentiate to PCF, *RDK1* mRNA levels decreased sharply within 24 hours in differentiating parasites (3.27-fold decrease compared to stumpy BSF) and further decreased in fully differentiated PCF parasites (4.15-fold decrease compared to stumpy BSF) (Fig. 1A). To determine relative RDK1 protein expression, we generated whole cell lysates from slender BSF and PCF parasites expressing RDK1 with a C-terminal 10xTy epitope tag from an endogenous locus and performed Western blot analysis (Fig. 1B). Contrary to the quantitative proteomics study that catalogued RDK1 as a PCF-upregulated protein, we observed that RDK1 expression was detectable in slender BSF parasites but decreased beneath the limit of detection in PCF parasites (Fig. 1B).

**Figure 1.**
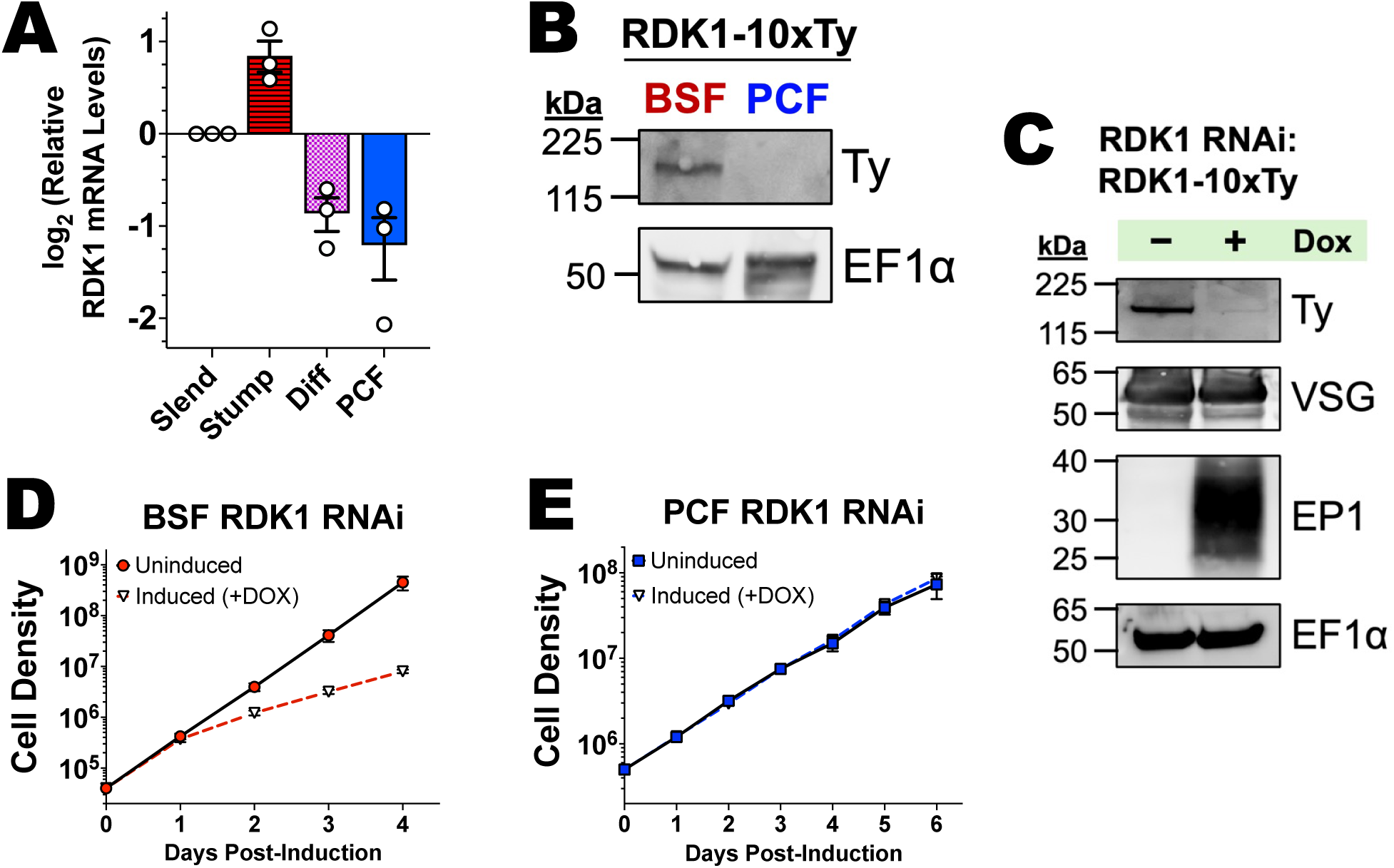
RDK1 is downregulated during BSF-to-PCF differentiation. (A) Quantitative RT-PCR analysis of relative *RDK1* mRNA levels in slender BSF (Slend), stumpy BSF (Stump), 24-hr differentiating (Diff), and PCF parasites normalized to an internal spike-in luciferase mRNA control. Slender BSF values were set to 1 (plotted on the y-axis as 0, as log_2_(1) = 0). Values were plotted and analyzed using GraphPad Prism 11.0. (B) Western blot of whole cell lysates of slender BSF (Slend), stumpy BSF (Stump), 24-hr and 48-hr differentiating (Diff), and PCF parasites to determine relative RDK1 protein levels. Anti-Ty was used to detect endogenously tagged RDK1. EF1α is used as a loading control. (C) Western blot of cell lysates of BSF uninduced (-Dox) and induced (+Dox) RDK1 RNAi parasites. Anti-Ty was used to detect RDK1 that was endogenously tagged with a C-terminal 10xTy epitope tag. VSG and EP1 are BSF and PCF surface markers, respectively. EF1α is used as a loading control. (D) Cell growth kinetics of BSF RDK1 RNAi parasites in the absence (uninduced) or presence (+Dox) of doxycycline for 4 days. Cell growth is plotted as cumulative cell density in log scale. (E) Cell growth kinetics of PCF RDK1 RNAi parasites in the absence (uninduced) or presence (+Dox) of doxycycline for 6 days. Cell growth is plotted as cumulative cell density in log scale.

We generated an inducible RDK1 RNAi cell line using BSF parasites expressing RDK1-10xTy from an endogenous locus to confirm substantial reduction of RDK1 protein following doxycycline treatment (Fig. 1C). Western blot analysis confirmed robust depletion of RDK1 in BSF RDK1 RNAi parasites that were treated with doxycycline for 48 hours. To confirm that our BSF RDK1 RNAi induced BSF-to-PCF differentiation in the EATRO1125 *T. brucei* strain, similar to the RDK1 RNAi cell line made with the 2T1 monomorphic BSF cell line in the initial RDK1 characterization study (17), we probed with life cycle markers for BSF parasites (VSG) and PCF parasites (EP1 procyclin). We observed that EP1 expression was induced by RDK1 RNAi; however, VSG expression was not dramatically decreased (Fig. 1C). This suggests that only a subset of parasites was expressing EP1 procyclin, similar to the original RDK1 RNAi cell line of the initial study. We also determined that cell growth rate of induced RDK1 RNAi parasites under standard BSF culture conditions were moderately reduced compared to uninduced controls (Fig. 1D). Although we observed that RDK1 protein expression was decreased in PCF parasites, there remained the possibility that RDK1 is expressed at low levels and remains important for PCF cell growth. To determine if RDK1 RNAi impacted PCF cell growth, we differentiated BSF RDK1 RNAi parasites into PCF parasites, quantified cumulative cell growth of PCF uninduced and induced RDK1 RNAi parasites and observed that RDK1 RNAi did not impact PCF cell growth (Fig. 1E). From these data, we conclude that RDK1 is expressed and essential for optimal cell growth of BSF parasites, and RDK1 is downregulated during BSF-to-PCF differentiation after which RDK1 is likely neither expressed nor essential in PCF parasites.

### Proximity-dependent protein labeling suggests dual localization in the cell membrane and internal mitochondrial subcompartments

The cellular localization of RDK1 has been unclear. Using a combination of a hypotonic swelling assay, a detergent solubilization assay, and immunofluorescence microscopy, the initial study that identified RDK1 concluded that RDK1 is a membrane-integrated protein that localized to the cell membrane and along the flagellum (17). However, a subsequent study that inducibly overexpressed RDK1 in BSF parasites concluded that RDK1 is localized to the endoplasmic reticulum (ER), as it co-localized with the ER protein marker BiP in their immunofluorescence microscopy images (24). We chose to utilize proximity-dependent protein biotinylation and identification (BioID) as an unbiased approach that did not rely on visual bias to determine cellular localization of RDK1 in BSF parasites. We generated a BSF cell line that fused the promiscuous biotin ligase (miniTurbo) to the C-terminal kinase domain of RDK1 in an endogenous locus (RDK1-miniTurbo). We harvested 5×10^8^ of parental BSF 90-13 and RDK1-miniTurbo parasites grown in fresh culture medium supplemented with 50 µM biotin for 12 hours. We then lysed parasites and incubated with streptavidin-conjugated magnetic beads followed by a series of stringent washes to remove non-specific proteins. Total input lysate and streptavidin affinity-enriched samples from parental 90-13 and RDK1-miniTurbo parasites were analyzed by Western blot to confirm that RDK1-miniTurbo was expressed and biotinylated proteins were enriched in the affinity-enriched samples in the RDK1-miniTurbo samples (Fig. 2A). Using streptavidin-conjugated horseradish peroxidase (HRP) as a probe, we observed robust enrichment of biotinylated proteins in the streptavidin-enriched samples of RDK1-miniTurbo parasites. We did not observe appreciable enrichment of biotinylated proteins in parental 90-13 parasites that do not express miniTurbo (Fig. 2A). Mass spectrometry identified 853 biotinylated proteins that were significantly enriched in RDK1-miniTurbo parasites compared to 90-13 parental controls, of which RDK1 itself is among the most enriched proteins (Fig. 2B, Table S1).

**Figure 2.**
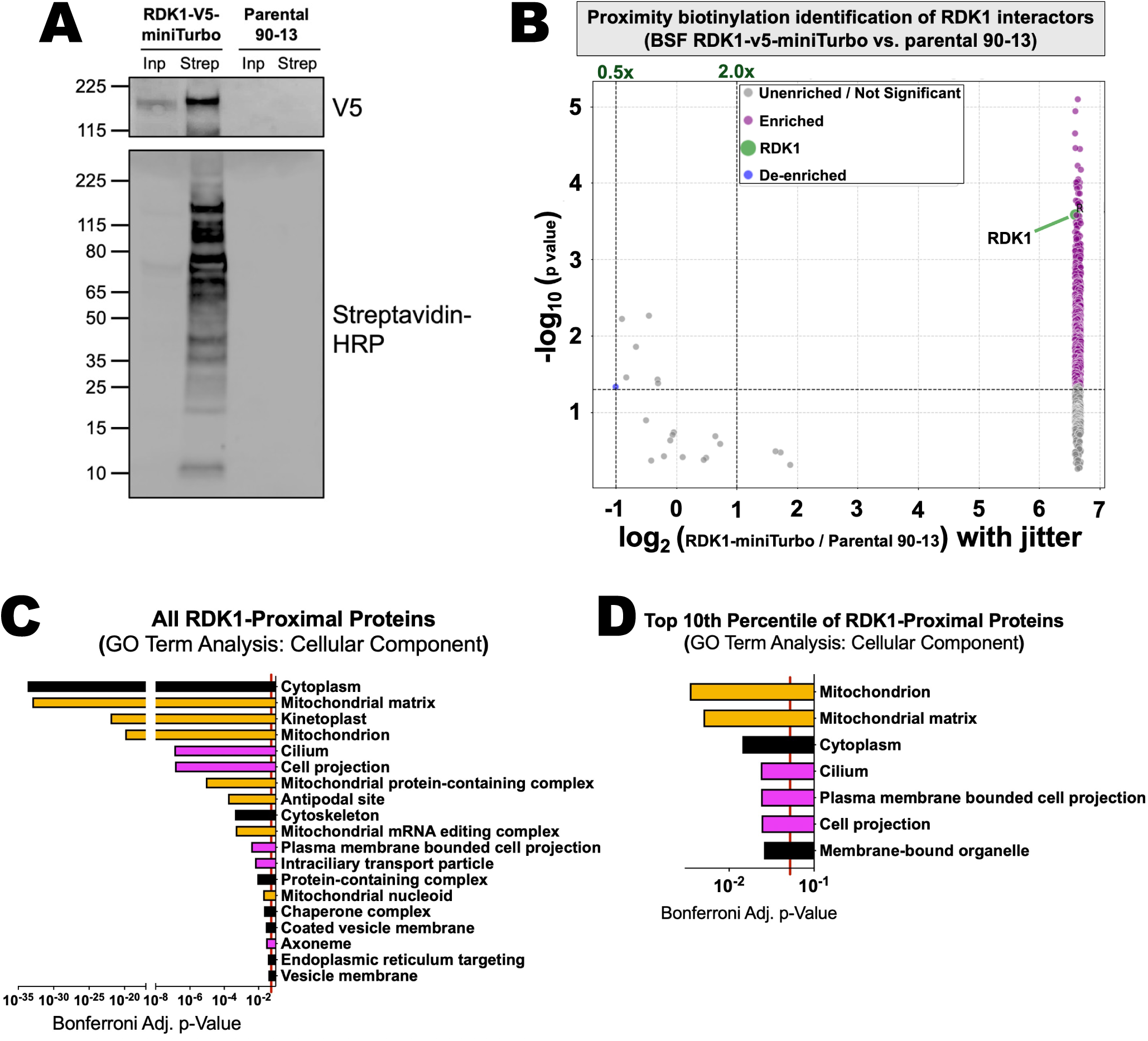
RDK1 has dual localization in the cell membrane and mitochondrial inner membrane. (A) Western blot analysis of total input (Inp) and streptavidin affinity-enriched (Strep) protein samples from BSF parental 90-13 and RDK1-miniTurbo lysates. Anti-V5 was used to visualize RDK1 endogenously tagged with a C-terminal V5-miniTurbo biotin ligase. Streptavidin-conjugated horseradish peroxidase (HRP). (B) Volcano plot of the 853 enriched biotinylated proteins identified by mass spectrometry in BSF RDK1-miniTurbo parasites compared to the parental BSF 90-13 control. (C-D) Gene Ontology Term analyses for cellular component for (C) all enriched RDK1-miniTurbo proximal interactions and (D) the top tenth percentile of RDK1-miniTurbo interactions. For panels C and D, yellow bars are associated with the mitochondrion and purple bars are associated with the cell membrane, cytoskeleton, or flagellum. The adjusted p-values with Bonferroni correction of p<0.05 were used as statistical significance cutoffs for ontology terms and are represented by a vertical red line. Redundant and obsolete terms were removed for clarity.

To deduce the unbiased localization of RDK1, we performed Gene Ontology (GO) Term analysis for cellular component of the 852 proximal proteins to RDK1 (Fig. 2C). We observed that the two most significant cellular component GO terms were the cytoplasm and the mitochondrial matrix (Fig. 2C). Considering that RDK1 is a membrane-integrated protein (17), these data suggest that RDK1 is both a dual-localized cell membrane protein and mitochondrial inner membrane protein with the C-terminal kinase domain of RDK1 oriented towards the cytoplasm and mitochondrial matrix, respectively. The mitochondrial localization is surprising and unexpected, as previous immunofluorescence studies did not have discrete staining patterns that would imply mitochondrial localization (17, 24). However, other statistically significant GO Terms for RDK1-proximal proteins include the mitochondrion, kinetoplast (mitochondrial DNA), the antipodal sites of the kinetoplast DNA, and the mitochondrial mRNA editing complex (Fig. 2C). Furthermore, GO Term analysis for cellular component for the top tenth percentile of RDK1-proximal proteins based on median enrichment values determined that the most enriched RDK1 interactions were mitochondrial and mitochondrial matrix proteins (Fig. 2D). In fact, the most enriched RDK1 interaction is the mitochondrial TCA cycle enzyme, alanine aminotransferase (Table 1). We also observed that several mitochondrial matrix heat shock proteins mtHsp10, mtHsp60, mtHsp70, and mtHsp84 are among the most enriched proteins (Table 1). Corroborating a mitochondrial localization, we also found that RDK1-miniTurbo comes in close proximity to mitochondrial protein import machinery, including the Atom69 mitochondrial receptor for the archaic translocase of the outer membrane (ATOM) complex (34) and a receptor of the translocase of the inner membrane (TIM) complex, TbTim62 (35)(36). This strongly implicates that RDK1 is natively imported into the mitochondrion. However, GO terms for the cytoplasm and the cilium (flagellum in *T. brucei*) were also statistically significant in the top tenth percentile of RDK1-proximal proteins (Fig. 2D). Thus, from these data, we conclude that RDK1 has a dual localization in the cell membrane and mitochondrial inner membrane in BSF.

**Table 1.**
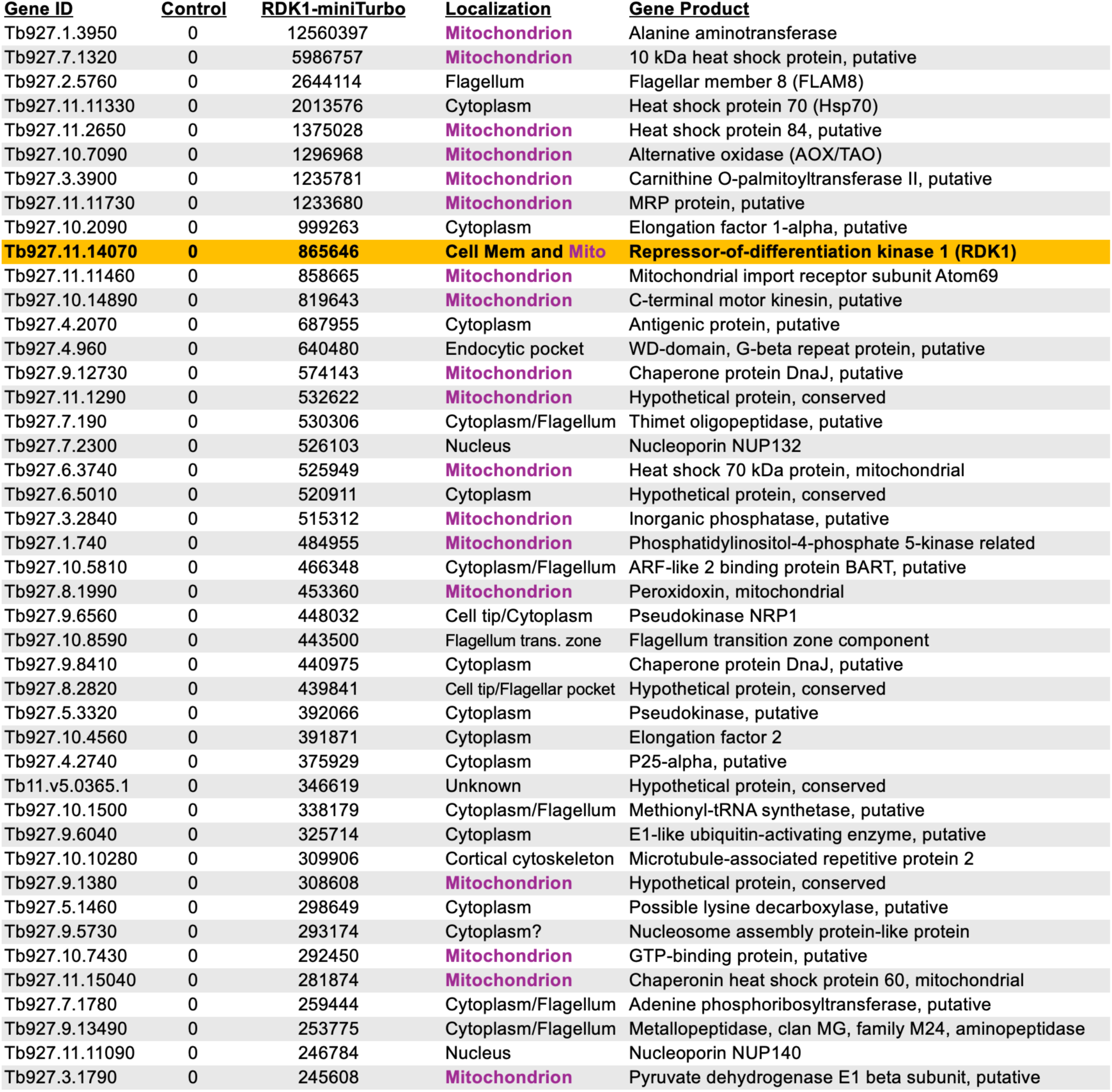
Top 5^th^ percentile of RDK1-proximal proteins in BSF parasites.

### RDK1 depletion upregulates expression of several mitochondrial proteins

To determine the impact of RDK1 depletion on the BSF proteome in the absence of differentiation cues, we harvested 5×10^8^ BSF uninduced and 48-hr induced RDK1 RNAi parasites grown in BSF HMI-9 medium under standard BSF culture conditions at 37°C. We hypotonically lysed the parasites in the presence of protease and phosphatase inhibitors before solubilization with 4% w/v SDS and 200 mM DTT. Mass spectrometry identification and quantitative global proteomics were performed to determine differentially upregulated (at least 1.50-fold increase) and downregulated (at lease 0.67-fold decrease) proteins in RDK1 RNAi parasites compared to uninduced control parasites (Fig. 3A). We observed that 19 identified proteins are significantly downregulated in RDK1 RNAi parasites, of which seven proteins are hypothetical proteins with unknown function (Table 2). RDK1 depletion also reduces protein levels of a serine-arginine protein kinase (SRPK1) and a cell division cycle phosphatase (CDC14), which play roles in stress response survival and cytokinesis, respectively (36, 37). Moreover, other downregulated proteins in RDK1 RNAi parasites are involved in cell cycle and cell division, including the cytokinesis initiation factor 2 (CIF2) and kinetoplast kinetochore proteins 2 and 7 (KKT2 and KKT7) (Table 2) (38–41). This corroborates the conclusion that RDK1 participates in the regulation of the cell cycle of BSF parasites.

**Figure 3.**
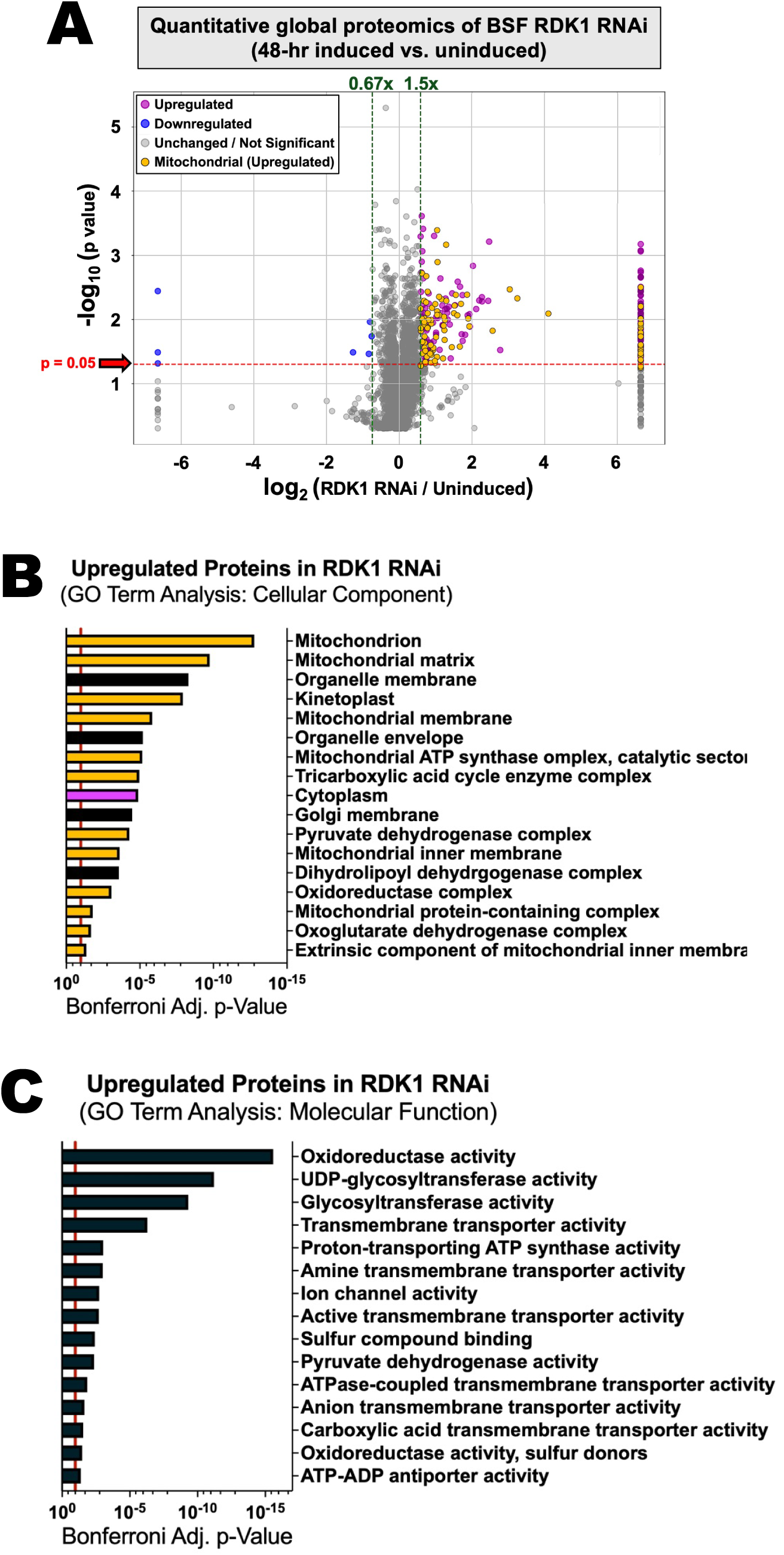
RDK1 depletion upregulates the abundance of mitochondrial proteins. (A) Volcano plot illustrating differential abundances of proteins in RDK1 RNAi parasites induced for 48 hours compared to the uninduced BSF control. Upregulated proteins (ý1.50x fold change) are denoted by purple dots; upregulated mitochondrial proteins are denoted by yellow dots; downregulated proteins (:s0.67x fold change) are denoted by blue dots; non-significant or non-regulated proteins are denoted by grey dots. (B-C) Gene Ontology Term analysis of upregulated proteins in RDK1 RNAi parasites that are detected in uninduced controls for (B) cellular component and (C) molecular function. For panels B-C, yellow bars are associated with the mitochondrion and purple bars are associated with the cell membrane, cytoskeleton, or flagellum. The adjusted p-values with Bonferroni correction of p<0.05 were used as statistical significance cutoffs for ontology terms and are represented by a vertical red line. Redundant and obsolete terms were removed for clarity.

**Table 2.** Downregulated proteins in BSF RDK1 RNAi parasites.

| **TriTrypDB**<br>**Gene ID** | **RDK1 RNAi vs.**<br>**Uninduced** | **Gene Product** |
| --- | --- | --- |
| Tb927.11.13490 | 0.01* | Hypothetical protein, conserved |
| Tb927.10.5080 | 0.01* | Hypothetical protein, conserved |
| Tb927.11.15080 | 0.01* | Transmembrane protein 14C, putative |
| Tb927.6.350 | 0.414 | Zinc finger protein, C2H2 type, putative |
| Tb927.1.2600 | 0.539 | Pumilio/PUF RNA-binding protein 9 (PUF9) |
| Tb927.11.12430 | 0.560 | Cell division cycle phosphatase 14 (CDC14), putative |
| Tb927.1.1740 | 0.573 | Microtubule-associated protein Futsch, putative |
| Tb927.4.4570 | 0.590 | Hypothetical protein, conserved |
| Tb927.9.14290 | 0.601 | Cytokinesis initiation factor 2 (CIF2) |
| Tb927.11.1030 | 0.605 | Kinetoplastid kinetochore protein 7 (KKT7) |
| Tb927.11.5740 | 0.615 | Formin-like protein |
| Tb927.1.4040 | 0.621 | Hypothetical protein, conserved |
| Tb927.7.960 | 0.627 | CMGC/SRPK protein kinase (SRPK1), putative |
| Tb927.1.4280 | 0.631 | Hypothetical protein, conserved |
| Tb927.1.1500 | 0.640 | Conserved protein, unknown function |
| Tb927.1.640 | 0.643 | Breast cancer gene 2 (BRCA2), putative |
| Tb927.5.2990 | 0.647 | Hypothetical protein, conserved |
| Tb927.11.10520 | 0.660 | Kinetoplastid kinetochore protein 2 (KKT2) |
| Tb927.7.7530 | 0.662 | Receptor-type adenylate cyclase GRESAG4, putative |

In contrast, we identified 253 upregulated proteins in RDK1 RNAi parasites (Fig. 3A, Table S2). Notably, 97 out of 253 upregulated proteins were mitochondrial proteins (Fig. 3A, yellow dots). Of the 253 upregulated, 81 proteins were detected only in the RDK1 RNAi parasites and not the uninduced BSF controls. The remaining 172 proteins were also detected in uninduced BSF parasites. GO Term analysis of upregulated proteins in RDK1 RNAi parasites confirmed that mitochondrial proteins comprise a significant fraction in upregulated proteins (Fig. 3B). Many of these mitochondrial proteins are cytochrome proteins, the first seven enzymes involved in the TCA cycle and proline catabolism (Tables 3 and 4, Table S2). Moreover, we also observe upregulation of some glycosomal metabolic proteins, including the glycosomal isoform of malate dehydrogenase and pyruvate phosphate dikinase (Tables 3 and 4, Table S2). In fact, GO Term analysis for molecular function confirm that there is a bias for metabolic processes including oxidoreductase activity, ATP synthase activity, pyruvate metabolism, and ADP/ATP transport (Fig. 3C). Thus, from these data, we conclude that RDK1 depletion initiates a mitochondrial proteomic remodeling that supports a shift towards amino acid catabolism in the absence of differentiation cues.

**Table 3.** Abbreviated list of the most abundant proteins detected only in RDK1 RNAi parasites.

| Gene ID<br>(from TriTrypDB) | Gene Annotation<br>(Protein Product) | Median<br>in RNAi |
| --- | --- | --- |
| Tb927.5.3630 | Glycosomal malate dehydrogenase | 608156 |
| Tb927.7.2640 | Cytoskeleton-associated protein 51 | 483473 |
| Tb927.7.5940 | Protein associated with differentiation 2 (PAD2) | 185654 |
| Tb927.9.6100 | PTP1-interacting protein of 39 kDa (PIP39) | 161771 |
| Tb927.11.6280 | Pyruvate phosphate dikinase (PPDK) | 154460 |
| Tb927.8.7600 | Amino acid transporter, putative | 138417 |
| Tb927.5.440 | Trans-sialidase/Concanavalin A-like lectin superfamily | 109533 |
| Tb927.9.12320 | Cystathione gamma lyase | 91569 |
| Tb927.11.16200 | Cytoskeleton-associated protein 17 | 73784 |
| Tb927.10.7630 | Plasma membrane choline transporter, putative | 59015 |
| Tb927.8.6170 | Transketolase | 47981 |
| Tb927.10.11970 | Glutamine aminotransferase (GlnAT) | 47946 |
| Tb11.02.5400 | Cystathionine beta-synthase, putative | 44761 |
| Tb927.10.6910 | S-Adenosyl-L-methionine-c-24-delta-sterol methyltransferase a | 44155 |
| Tb927.4.3500 | Amastin surface glycoprotein, putative | 43483 |
| Tb927.11.2410 | Flabarin, putative | 42091 |
| Tb927.10.9080 | Pteridine transporter, putative | 39367 |
| Tb927.11.12830 | Acyl-CoA-binding protein, putative | 38485 |
| Tb927.10.7290 | Flagellar member 2 (FLAM2) | 37779 |
| Tb927.8.3620 | Folate transporter 1 (FT1) | 37484 |
| Tb927.10.8490 | Hexose transporter (THT1) | 33793 |
| Tb927.2.5530 | Present in the outer mitochondrial membrane proteome 22-1 | 33497 |
| Tb927.8.6010 | Heme response-1 protein | 33155 |
| Tb927.10.11220 | Procyclic form surface phosphoprotein | 32125 |
| Tb927.7.4390 | Threonine synthase, putative | 30683 |
| Tb927.3.4500 | Fumarate hydratase class I, cytosolic | 29872 |
| Tb927.11.4680 | SURF1 family protein, putative | 27843 |
| Tb927.5.1060 | Mitochondrial processing peptidase, beta subunit | 27577 |
| Tb927.9.7470 | Purine nucleoside transporter | 26948 |
| Tb927.10.4330 | Branched chain alpha-ketoacid dehydrogenase E1-beta subunit | 26109 |
| Tb927.7.7210 | Present in the outer mitochondrial membrane proteome 37 | 25891 |
| Tb927.11.3270 | Squalene monooxygenase, putative | 24827 |
| Tb927.10.6190 | Aldehyde dehydrogenase, putative | 24397 |
| Tb927.9.3170 | Cytochrome oxidase subunit V (COV/CoxV) | 24360 |
| Tb927.9.4310 | Tricarboxylate carrier, putative | 24179 |
| Tb927.11.8990 | Cation transporter, putative | 22789 |
| Tb927.5.320 | Adenylate and guanylate cyclase domain-containing protein (GRESAG4) | 21751 |
| Tb927.8.2970 | Hypothetical protein, conserved | 21657 |
| Tb927.7.2680 | Zinc finger protein family member (ZC3H22) | 21070 |
| Tb927.11.7040 | Pterin-4-alpha-carbinolamine dehydratase, putative | 20654 |
| Tb927.11.3980 | Metallopeptidase, Clan ME, Family 16 | 17854 |
| Tb927.11.16220 | Cytochrome b-c1 complex subunit, putative | 17165 |
| Tb927.5.2160 | Domain of unknown function (DUF306), putative | 17119 |
| Tb927.8.1610 | Major surface protease gp63, putative | 16115 |
| Tb927.11.16750 | Hypothetical protein, conserved | 14893 |
| Tb927.10.13430 | Citrate synthase, putative | 14856 |
| Tb927.11.14610 | Procyclin-associated gene 4 (PAG4) | 14826 |
| Tb927.10.5540 | Hypothetical protein, conserved | 14207 |
| Tb927.11.3750 | NADH-cytochrome b5 reductase, putative | 13385 |
| Tb927.1.4100 | Cytochrome oxidase subunit IV (COIV/CoxIV) | 13329 |
| Tb927.7.2670 | Zinc finger protein family member (ZC3H21) | 13176 |
| Tb927.9.2320 | Methyltransferase domain-containing protein, putative | 12681 |
| Tb927.6.2390 | Hypothetical protein, conserved | 12222 |
| Tb927.9.2450 | Electron transport protein SCO1/SCO2, putative | 11919 |

**Table 4.**
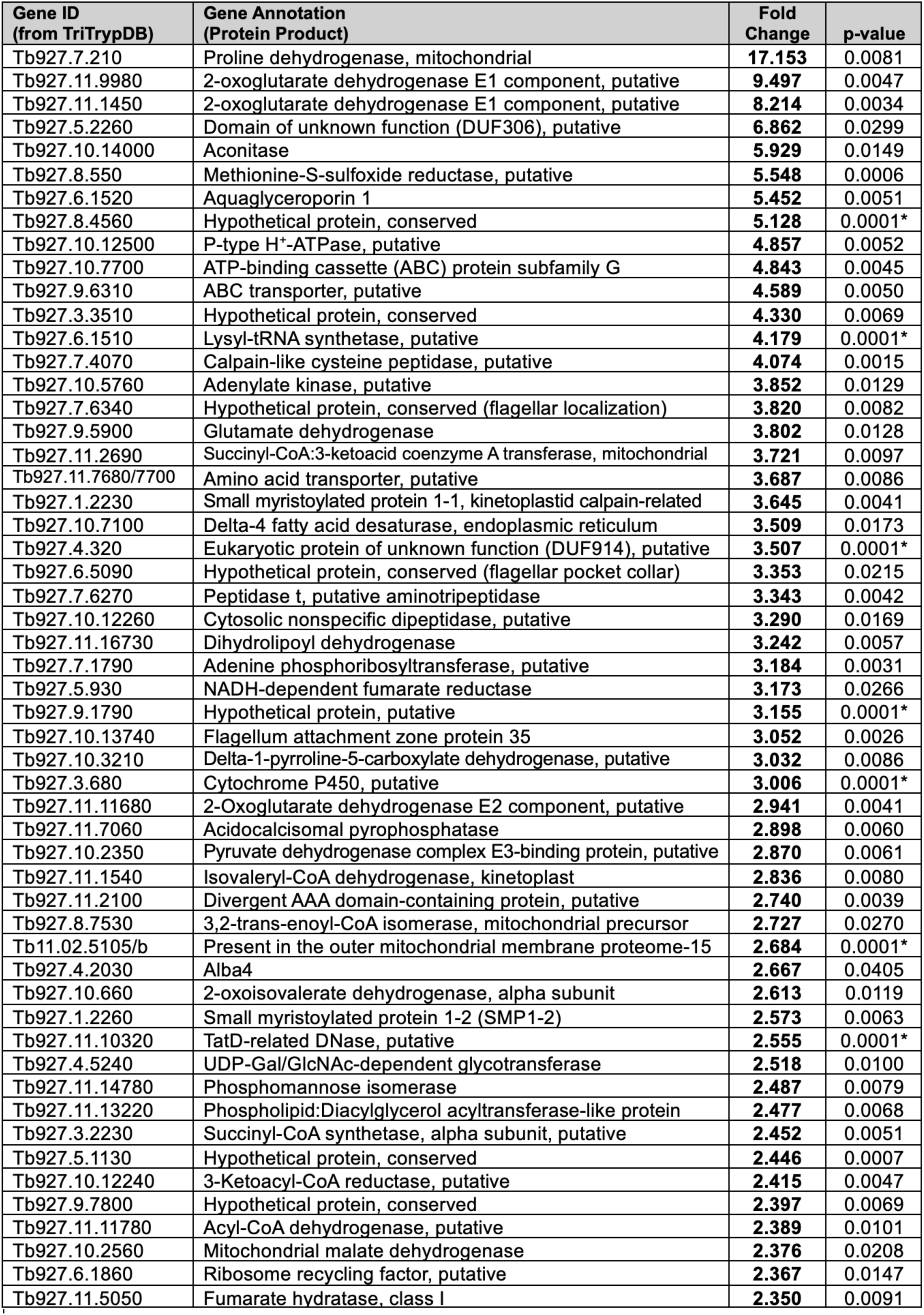
Abbreviated list of upregulated proteins in BSF RDK1 RNAi parasites.

| **Gene ID**<br>**(from TriTrypDB)** | **Gene Annotation**<br>**(Protein Product)** | **Fold**<br>**Change** | **p-value** |
| --- | --- | --- | --- |
| Tb927.7.210 | Proline dehydrogenase, mitochondrial | **17.153** | 0.0081 |
| Tb927.11.9980 | 2-oxoglutarate dehydrogenase E1 component, putative | **9.497** | 0.0047 |
| Tb927.11.1450 | 2-oxoglutarate dehydrogenase E1 component, putative | **8.214** | 0.0034 |
| Tb927.5.2260 | Domain of unknown function (DUF306), putative | **6.862** | 0.0299 |
| Tb927.10.14000 | Aconitase | **5.929** | 0.0149 |
| Tb927.8.550 | Methionine-S-sulfoxide reductase, putative | **5.548** | 0.0006 |
| Tb927.6.1520 | Aquaglyceroporin 1 | **5.452** | 0.0051 |
| Tb927.8.4560 | Hypothetical protein, conserved | **5.128** | 0.0001* |
| Tb927.10.12500 | P-type H<sup>+</sup>-ATPase, putative | **4.857** | 0.0052 |
| Tb927.10.7700 | ATP-binding cassette (ABC) protein subfamily G | **4.843** | 0.0045 |
| Tb927.9.6310 | ABC transporter, putative | **4.589** | 0.0050 |
| Tb927.3.3510 | Hypothetical protein, conserved | **4.330** | 0.0069 |
| Tb927.6.1510 | Lysyl-tRNA synthetase, putative | **4.179** | 0.0001* |
| Tb927.7.4070 | Calpain-like cysteine peptidase, putative | **4.074** | 0.0015 |
| Tb927.10.5760 | Adenylate kinase, putative | **3.852** | 0.0129 |
| Tb927.7.6340 | Hypothetical protein, conserved (flagellar localization) | **3.820** | 0.0082 |
| Tb927.9.5900 | Glutamate dehydrogenase | **3.802** | 0.0128 |
| Tb927.11.2690 | Succinyl-CoA:3-ketoacid coenzyme A transferase, mitochondrial | **3.721** | 0.0097 |
| Tb927.11.7680/7700 | Amino acid transporter, putative | **3.687** | 0.0086 |
| Tb927.1.2230 | Small myristoylated protein 1-1, kinetoplastid calpain-related | **3.645** | 0.0041 |
| Tb927.10.7100 | Delta-4 fatty acid desaturase, endoplasmic reticulum | **3.509** | 0.0173 |
| Tb927.4.320 | Eukaryotic protein of unknown function (DUF914), putative | **3.507** | 0.0001* |
| Tb927.6.5090 | Hypothetical protein, conserved (flagellar pocket collar) | **3.353** | 0.0215 |
| Tb927.7.6270 | Peptidase t, putative aminotripeptidase | **3.343** | 0.0042 |
| Tb927.10.12260 | Cytosolic nonspecific dipeptidase, putative | **3.290** | 0.0169 |
| Tb927.11.16730 | Dihydrolipoyl dehydrogenase | **3.242** | 0.0057 |
| Tb927.7.1790 | Adenine phosphoribosyltransferase, putative | **3.184** | 0.0031 |
| Tb927.5.930 | NADH-dependent fumarate reductase | **3.173** | 0.0266 |
| Tb927.9.1790 | Hypothetical protein, putative | **3.155** | 0.0001* |
| Tb927.10.13740 | Flagellum attachment zone protein 35 | **3.052** | 0.0026 |
| Tb927.10.3210 | Delta-1-pyrroline-5-carboxylate dehydrogenase, putative | **3.032** | 0.0086 |
| Tb927.3.680 | Cytochrome P450, putative | **3.006** | 0.0001* |
| Tb927.11.11680 | 2-Oxoglutarate dehydrogenase E2 component, putative | **2.941** | 0.0041 |
| Tb927.11.7060 | Acidocalcisomal pyrophosphatase | **2.898** | 0.0060 |
| Tb927.10.2350 | Pyruvate dehydrogenase complex E3-binding protein, putative | **2.870** | 0.0061 |
| Tb927.11.1540 | Isovaleryl-CoA dehydrogenase, kinetoplast | **2.836** | 0.0080 |
| Tb927.11.2100 | Divergent AAA domain-containing protein, putative | **2.740** | 0.0039 |
| Tb927.8.7530 | 3,2-trans-enoyl-CoA isomerase, mitochondrial precursor | **2.727** | 0.0270 |
| Tb11.02.5105/b | Present in the outer mitochondrial membrane proteome-15 | **2.684** | 0.0001* |
| Tb927.4.2030 | Alba4 | **2.667** | 0.0405 |
| Tb927.10.660 | 2-oxoisovalerate dehydrogenase, alpha subunit | **2.613** | 0.0119 |
| Tb927.1.2260 | Small myristoylated protein 1-2 (SMP1-2) | **2.573** | 0.0063 |
| Tb927.11.10320 | TatD-related DNase, putative | **2.555** | 0.0001* |
| Tb927.4.5240 | UDP-Gal/GlcNAc-dependent glycotransferase | **2.518** | 0.0100 |
| Tb927.11.14780 | Phosphomannose isomerase | **2.487** | 0.0079 |
| Tb927.11.13220 | Phospholipid:Diacylglycerol acyltransferase-like protein | **2.477** | 0.0068 |
| Tb927.3.2230 | Succinyl-CoA synthetase, alpha subunit, putative | **2.452** | 0.0051 |
| Tb927.5.1130 | Hypothetical protein, conserved | **2.446** | 0.0007 |
| Tb927.10.12240 | 3-Ketoacyl-CoA reductase, putative | **2.415** | 0.0047 |
| Tb927.9.7800 | Hypothetical protein, conserved | **2.397** | 0.0069 |
| Tb927.11.11780 | Acyl-CoA dehydrogenase, putative | **2.389** | 0.0101 |
| Tb927.10.2560 | Mitochondrial malate dehydrogenase | **2.376** | 0.0208 |
| Tb927.6.1860 | Ribosome recycling factor, putative | **2.367** | 0.0147 |
| Tb927.11.5050 | Fumarate hydratase, class I | **2.350** | 0.0091 |

### RDK1 depletion does not significantly impact mitochondrial physiologic state indicators of electron transport chain activity

Our global proteomics data indicate that RDK1 depletion causes BSF parasites to partially remodel the mitochondrial proteome with the upregulation of several cytochrome proteins and other metabolic enzymes. Therefore, we wanted to determine if RDK1 depletion altered other physiological indicators of mitochondrial function to a PCF-like state. First, we quantified mitochondrial superoxide levels of induced RDK1 RNAi parasites compared both to uninduced BSF and wildtype PCF parasites. The PCF mitochondrion produces reactive oxygen species as a biological byproduct of electron transport chain activity (42, 43). We stained parasites with MitoSOX Superoxide Indicator dye, which is converted to a fluorescent molecule when it reacts with the superoxide radical, and measured fluorescence by flow cytometry. We observe that RDK1 depletion causes no significant increase in mitochondrial superoxide levels compared to uninduced controls (Fig. 4A). As controls, we observed that PCF mitochondrial superoxide levels are 2.87-fold higher than uninduced BSF parasites. Furthermore, pre-treatment of PCF parasites with 100 µM paraquat for 30 minutes before MitoSOX staining was able to further increase mitochondrial superoxide levels by 36% compared to untreated PCF parasites (Fig. 4A). We measured mitochondrial membrane potential of RDK1 RNAi parasites compared to uninduced BSF and wildtype PCF controls. The mitochondrial membrane potential of PCF parasites is elevated compared to BSF due to the proton-pumping activity of the electron transport chain in the active PCF mitochondrion. We stained uninduced BSF, 48-hr induced RDK1 RNAi, and wildtype PCF parasites with TMRE and measured fluorescence by flow cytometry (Fig. 4B). RDK1 depletion caused a 1.40-fold increase in mitochondrial membrane potential, which was intermediate between uninduced BSF and wildtype PCF parasites. As a control, we pre-treated PCF parasites with 10 µM FCCP for 10 minutes, which caused a near-complete collapse of mitochondrial membrane potential (Fig. 4B). Taken together, these data suggest that the proteomic remodeling of the BSF mitochondrion caused by RDK1 depletion does not significantly impact physiologic indicators of electron transport chain activity.

**Figure 4.**
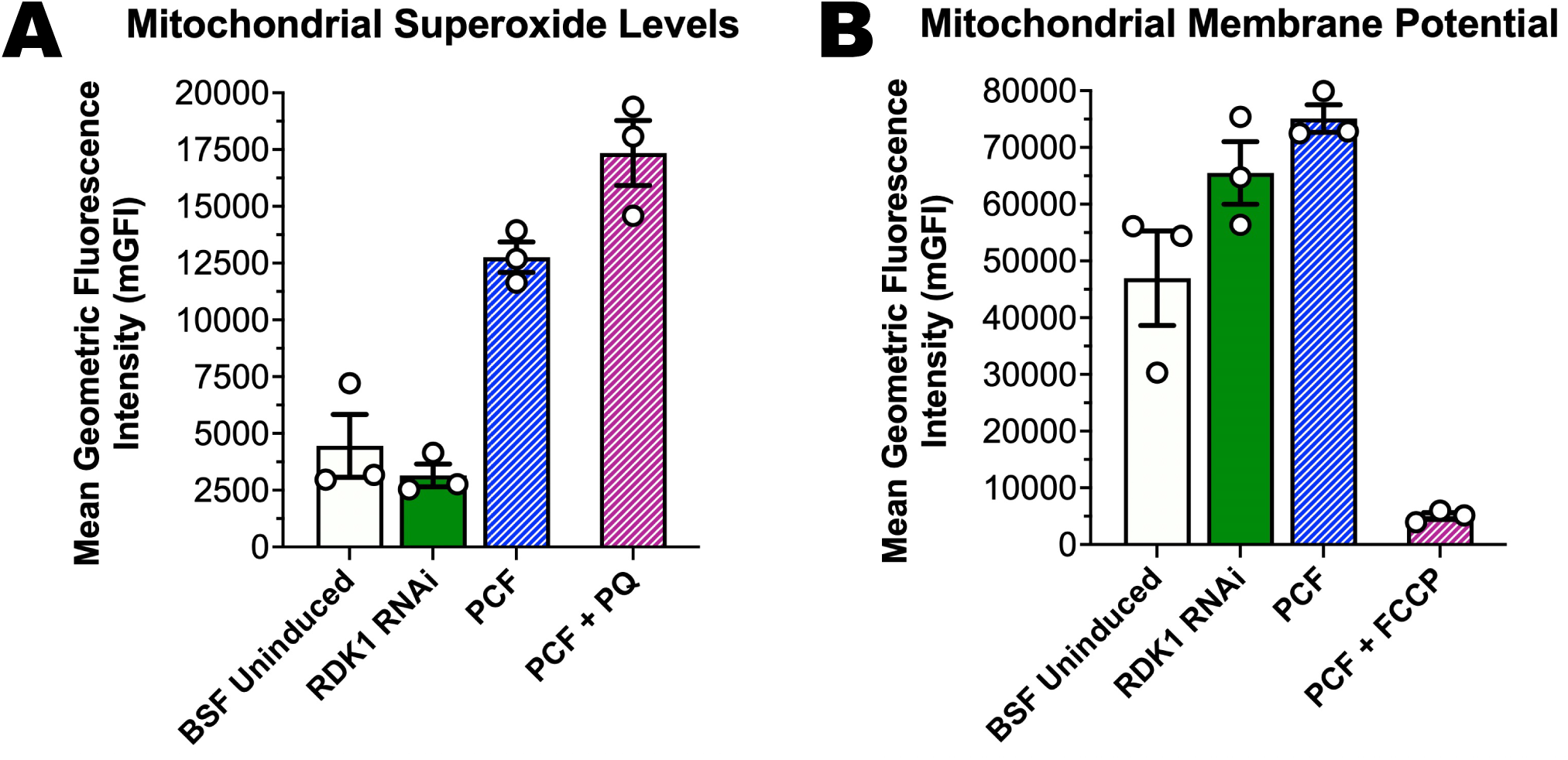
Mitochondrial physiologic state indicators of electron transport chain activity are mostly unaffected in RDK1 RNAi parasites. (A) Quantification of relative levels of mitochondrial superoxide by MitoSOX™ staining in BSF uninduced, 48-hr induced RDK1 RNAi, and PCF wildtype parasites. PCF parasites pre-treated with 100 µM paraquat (PQ) were included as a control for the induction of superoxide. (B) Quantification of mitochondrial membrane potential by TMRE staining in BSF uninduced, 48-hr induced RDK1 RNAi, and PCF wildtype parasites. PCF parasites were treated with 10 µM FCCP as a control for membrane depolarization. The values in panels B-C were plotted and analyzed using GraphPad Prism 11.0.

### RDK1 depletion dysregulates phosphorylation events near the cell membrane and the flagellum

The newfound discovery that RDK1 is proximal to many mitochondrial proteins combined with the pronounced impact on the mitochondrial proteome upon RDK1 depletion led us to hypothesize that RDK1 regulates the mitochondrial phosphoproteome. Phosphopeptide enrichment and mass spectrometry analysis of 48-hr induced RDK1 RNAi parasites compared to uninduced BSF parasites identified 339 phosphopeptides that were increased at least 2.0-fold that mapped 144 proteins and identified 2,114 phosphopeptides that were decreased at least 0.5-fold that mapped to 399 proteins (Fig. 5A, Table S3). In total, 527 proteins displayed differentially phosphorylated phosphosites in RDK1 RNAi parasites, which means only 16 proteins have phosphorylation sites that were hyperphosphorylated while other phosphorylation sites on the same protein were hypophosphorylated.

**Figure 5.**
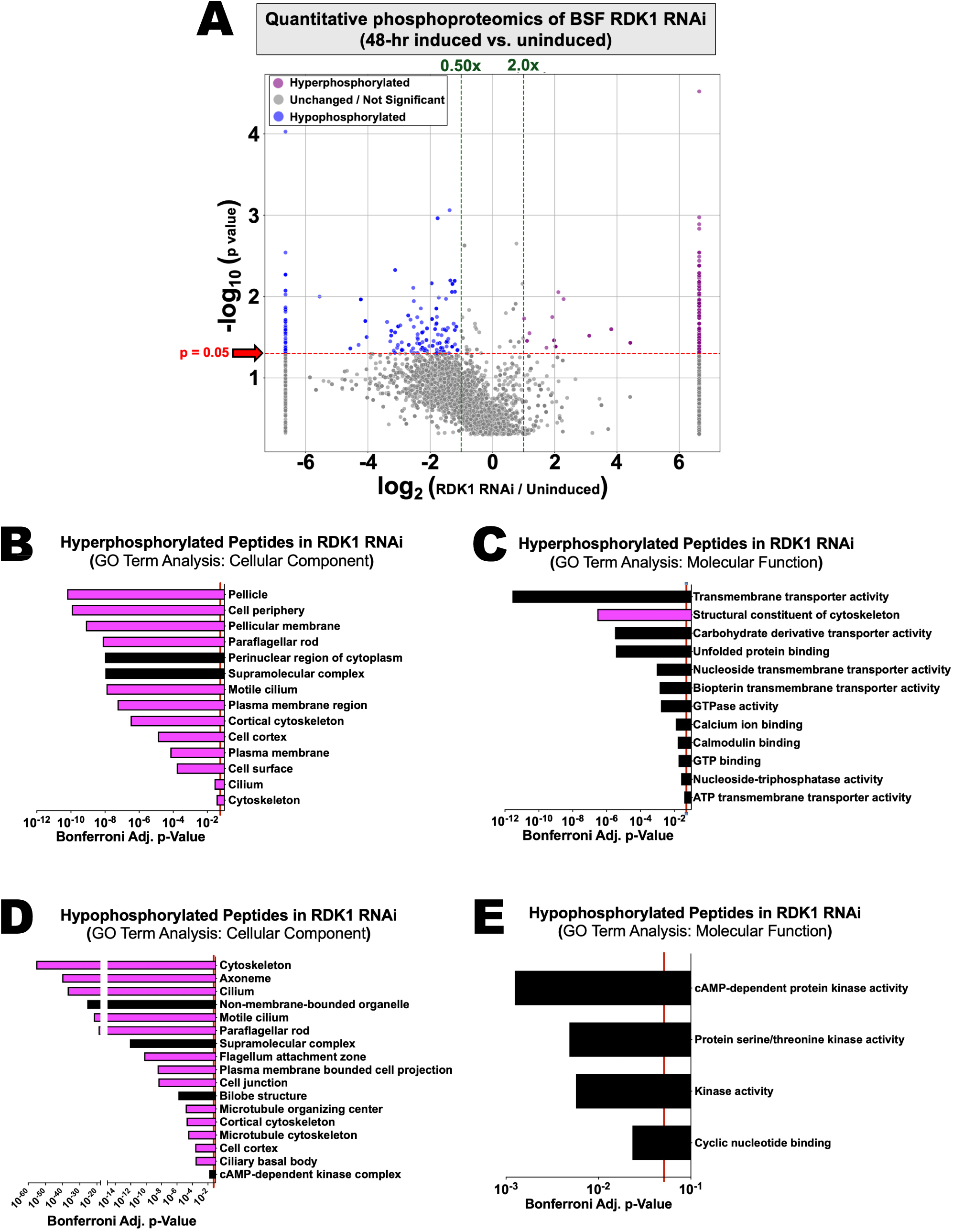
RDK1 depletion affects the phosphoproteome near the cell membrane and flagellum. (A) Volcano plot illustrating differential abundances of phosphopeptides in RDK1 RNAi parasites induced for 48 hours compared to the uninduced BSF control. Hyperphosphorylated peptides (ý2.00x fold change) are denoted by purple dots; hypophosphorylated peptides (:s0.50x fold change) are denoted by blue dots; non-significant or non-regulated phosphopeptides are denoted by grey dots. (B-C) Gene Ontology Term analysis of hyperphosphorylated peptides in RDK1 RNAi parasites compared to uninduced controls for (B) cellular component and (C) molecular function. (D-E) Gene Ontology Term analysis of hypophosphorylated peptides in RDK1 RNAi parasites compared to uninduced controls for (D) cellular component and (E) molecular function. For panels B-E, yellow bars are associated with the mitochondrion and purple bars are associated with the cell membrane, cytoskeleton, or flagellum. The adjusted p-values with Bonferroni correction of p<0.05 were used as statistical significance cutoffs for ontology terms and are represented by a vertical red line. Redundant and obsolete terms were removed for clarity.

Although the depletion of RDK1 favored a dephosphorylation trend, there were increases in the phosphorylation of 144 proteins. GO Term analysis indicates that hyperphosphorylated proteins are generally associated with pellicular array of microtubules, pellicular membrane, flagellum, paraflagellar rod, and plasma membrane (Fig. 5B). GO Term analysis also suggests that hyperphosphorylated proteins in RDK1 RNAi are generally involved in transmembrane transport of molecules, cytoskeletal structure, calcium binding, and GTP binding and hydrolysis (Fig. 5C). This suggests that RDK1 may potentially inhibit the activity of other protein kinases that target cytoskeletal and pellicle structures to control cell morphology and cytokinesis.

GO Term analysis of the 399 hypophosphorylated proteins reveals that RDK1-dependent phosphorylation events are concentrated in the cytoskeleton, flagellum, paraflagellar rod, and the axoneme (Fig. 5D). Molecular function of hypophosphorylated proteins largely correlate with protein kinase activity, cyclic nucleotide binding, and cAMP-dependent protein kinase activity (Fig. 5E). Most of the hypophosphorylated kinases in RDK1 RNAi parasites are putative, uncharacterized kinases. However, RDK1 depletion also affects the phosphorylation of cdc2-related kinase 9 (CRK9), which regulates mRNA *trans*-splicing and the phosphorylation of RNA polymerase II subunit RPB1 (44). RDK1 depletion also impacts the phosphorylation of a polo-like kinase orthologue, which functions in regulating the cell cycle (45). Interestingly, serine-392 of RDK2 was hypophosphorylated in RDK1 RNAi parasites (10.00-fold less phosphorylation than uninduced BSF parasites); although, serine-392 hypophosphorylation likely does not completely negate RDK2 function, as RDK2 RNAi parasites have distinct phenotypes from RDK1 RNAi parasites (17). Surprisingly, however, there was no significant enrichment of mitochondrial proteins that were either hypophosphorylated or hyperphosphorylated, and RDK1-dependent phosphoregulation is largely spatially restricted near the cell membrane, flagellum, and cytoskeleton. This implies that, despite frequent association with mitochondrial proteins (Fig. 2C-2D), RDK1 does not display robust kinase activity in the mitochondrion. From these data, we conclude that RDK1 regulates the phosphoproteome in its cell membrane localization and that RDK1 impacts both phosphorylation signaling and cyclic nucleotide signaling.

### RDK1 possesses kinase activity but not adenyl cyclase activity

Hypophosphorylated proteins in RDK1 RNAi parasites are associated with functions both in kinase signaling and cyclic nucleotide signaling, particularly cyclic AMP (cAMP) signaling. Notably, RDK1 has two predicted enzymatic domains: the C-terminal kinase domain and a central adenyl or guanyl cyclase domain (Fig. 6A) (24). We wanted to determine if RDK1 has both kinase and adenyl cyclase activities. To quantify kinase activity, we lysed BSF RDK1-10xTy parasites under non-denaturing conditions, and we immunoprecipitated (IP) RDK1 with anti-Ty antibody-conjugated protein A sepharose beads. RDK1 IP samples were then used in the ADP-Glo Kinase Assay (Promega) and kinase activity was measured by luminometry (Fig. 6B). Parental BSF 90-13 parasites that do not express a Ty-tagged protein were used as negative control. BSF parasites that endogenously express C-terminally 10xTy-tagged MEKK1, which was previously determined to have kinase activity, were used as a positive control. Our data indicate that RDK1 has robust kinase activity, similar to MEKK1 (24) (Fig. 6B). To determine adenyl cyclase activity, we performed RDK1 IP in the same manner as the kinase assays and used RDK1 IP samples with the cAMP-Glo Assay (Promega) (Fig. 6C). Our data suggest that RDK1 does not possess adenyl cyclase activity. It is possible that our *in vitro* adenyl cyclase activity assay is not optimal for RDK1. Thus, we determined if RDK1 depletion impacts intracellular cAMP levels using a previously established protocol (46). To determine relative intracellular cAMP levels, we lysed uninduced and 48-hr induced RDK1 RNAi parasites in induction buffer containing phosphodiesterase inhibitors. We observed that RDK1 depletion does not significantly cAMP levels (Fig. 6D). From these data, we conclude that RDK1 possesses kinase activity, despite the lack of phosphoregulation in the mitochondrion, but does not possess adenyl cyclase activity. We speculate that cyclic nucleotide signaling is likely impacted through the phosphorylation of proteins involved in cAMP signaling.

**Figure 6.**
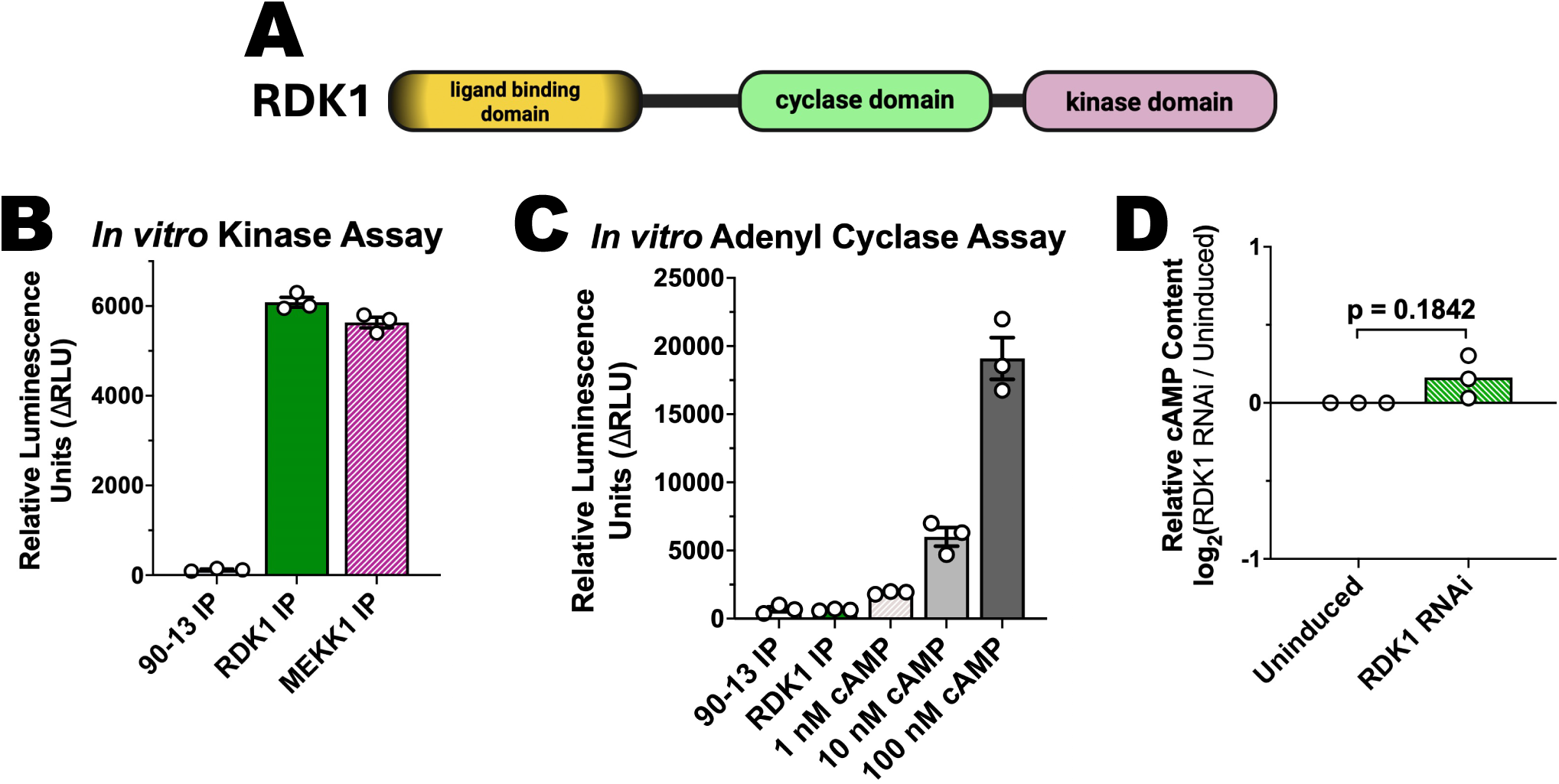
RDK1 possesses kinase activity but not adenyl cyclase activity *in vitro*. (A) Schematic of RDK1 domain architecture. (B) *In vitro* kinase activity assays using Ty-immunoprecipitated (IP) samples from BSF parental 90-13, RDK1-10xTy, and MEKK1-10xTy parasites. A buffer blank with no IP samples added was used as a negative background control. (C) *In vitro* adenyl cyclase activity assays using Ty-immunoprecipitated (IP) samples from BSF parental 90-13 and RDK1-10xTy parasites. A buffer blank with no cAMP added (0 nM) was used as a negative background control. Positive controls for cAMP-induced protein kinase A activity have 1 nM, 10 nM, or 100 nM of cAMP added. (D) Measurements of intracellular cAMP levels in 48-hr induced RDK1 RNAi parasites compared to uninduced controls. The values in panels B-D were plotted and analyzed using GraphPad Prism 11.0.

## DISCUSSION

The initial study that characterized RDK1 used the monomorphic 2T1 BSF cell line, which lacks the ability to differentiate into stumpy BSF due to defunct, yet unidentified, signaling pathways. This makes the 2T1 cell line suboptimal for differentiation studies that involve cell signaling. Our study utilized the pleomorphic EATRO1125 strain that retains the abilities to readily differentiate into stumpy BSF and PCF parasites. Our study represents the first systems-level characterization of RDK1 in *T. brucei* that emphasized the foundational analysis of RDK1 protein interactions and the effect of RDK1 depletion on the BSF proteome and phosphoproteome instead of overall gene expression. Our work revealed that RDK1, when expressed from an endogenous locus, localizes the cell membrane and the mitochondrial inner membrane. This was unexpected, as previous immunofluorescence experiments did not display clear mitochondrial staining (17, 24). However, some images in the initial characterization study had possible RDK1 signal near the kinetoplast (17), which coincides with our TurboID data that enriches several kinetoplast proteins. Another study implicated RDK1 was involved in repressing the uridine-insertion (U-indel) editing of the mitochondrial-encoded *COII* mRNA (30). While this observation was initially assumed to be indirect because RDK1 was not expected to be a mitochondrial protein, as it was previously detected in mitochondrial proteomes (47). However, RDK1 may not be easily detected by mass spectrometry without prior enrichment as it is often missing in several global proteomics datasets, including our global proteomics dataset of RDK1 RNAi parasites (Table S2). Moreover, our data shows that RDK1 is proximal to 19 proteins that function in U-indel mRNA editing and includes proteins that are assembled in the RNA editing catalytic complexes (RECC), the RNA editing substrate-binding complex (RESC), and crucial accessory proteins. Thus, this opens the possibility that RDK1 may directly impact U-indel editing of *COII* mRNA.

RDK1 depletion causes BSF parasites to initiate a differentiation program reflected in the proteome. Our data demonstrate that many mitochondrial and glycosomal metabolic enzymes are upregulated in RDK1 RNAi parasites in the absence of differentiation cues such as temperature reduction, *cis*-aconitate, glucose deprivation, or a change in culture medium. However, in the absence of these differentiation cues, we observe a lack of physiologic indicators that the parasite mitochondrion has resumed oxidative phosphorylation. Thus, the RDK1-depleted BSF mitochondrion likely does not perform oxidative phosphorylation; however, it may be primed for increased ATP production by substrate-level phosphorylation as TCA enzymes are increased, which the BSF mitochondrion can perform (48). Furthermore, there is an increase in glucose transporters in the cell membrane and some glycolytic enzymes, which suggests a potential increase in ATP production by glycolysis. However, future studies will have to be performed to determine if the RDK1-depleted mitochondrion produces more ATP than uninduced controls. Thus, our study suggests that RDK1 depletion may prime BSF parasites for metabolic reprogramming towards a PCF-like metabolism.

The initial RDK1 study published in 2014 suggested that RDK1 is likely a promoter of stumpy BSF rather than an outright repressor of PCF differentiation. Our data agrees with this speculation in that there is no clear functional remodeling of the mitochondrion; however, this assumes that RDK1 is a constitutive repressor. While the initial study from the Mottram lab and our study both use RNAi to deplete RDK1, it must be noted that RDK1 likely remains expressed in wildtype BSF parasites early during BSF-to-PCF differentiation. Our data demonstrate that RDK1 is downregulated in PCF (Fig. 1A-1B); however, RDK1 activity may be modified or circumvented in early in differentiation-stimulated parasites before its eventual downregulation in PCF parasites.

RDK1 has both predicted kinase and adenyl/guanyl cyclase domains. Although we confirm kinase activity, we do not detect adenyl cyclase activity of RDK1 *in vitro*, and we do not observe an impact on cAMP levels after RDK1 depletion. It is possible that the dysregulated phosphorylation of several cyclic nucleotide signaling proteins may have increased and decreased function that ultimately buffer impacts on cAMP levels. However, it is also possible that RDK1 functions as a guanyl cyclase and not an adenyl cyclase. In fact, the domain architecture of RDK1 resembles receptor guanyl cyclases in metazoans, which typically have an N-terminal ligand-binding domain, a central kinase homology domain, and a C-terminal guanyl cyclase domain. While receptor guanyl cyclases often do not exhibit *in vivo* kinase activity, the orientation of the cyclase and kinase domains of RDK1 are inverted, which may allow permissive access of the RDK1 kinase domain to *in vivo* substrates. However, we demonstrate that the phosphorylation status of mitochondrial proteins is largely unaffected, despite the clear proximity of mitochondrial proteins with the kinase domain of RDK1. This suggests that the kinase domain of RDK1 is likely inactivated in the mitochondrion. Interestingly, enzymatic activity of receptor guanyl cyclases can be regulated through the association of heat shock proteins and the kinase homology domain (25, 49). Our data show that RDK1 is proximal to several mitochondrial heat shock proteins. Thus, it is possible that the association of mitochondrial heat shock proteins in the mitochondrial matrix masks kinase activity. However, it remains unknown if this association changes during early BSF-to-PCF differentiation when the temperature decreases.

In conclusion, we assert that RDK1 dual-localizes to the cell membrane and mitochondrial inner membrane and has localization-specific kinase activity. Depletion of RDK1 dysregulates the phosphoproteome in the cell membrane locale and may impact cell division and cell morphology, but RDK1 may exert kinase-independent functions to maintain the mitochondrial proteome. Lastly, circumvention of RDK1 function during BSF-to-PCF differentiation, either through downregulation or post-translational mechanisms, may prime parasites for metabolic reprogramming as *T. brucei* parasites must transition from glucose dependence to amino acid dependence as a primary means for ATP production. Further studies are warranted to decipher kinase-dependent functions from kinase-independent functions of RDK1.

## MATERIALS AND METHODS

### Cultivation of trypanosomes and generation of transgenic parasites

In this study, we utilized the pleomorphic EATRO1125 AnTat1.1 *T. brucei* 90-13 cell line that was genetically engineered to constitutively express the T7 RNA polymerase and TetR protein for tetracycline-inducible gene expression (50). Slender bloodstream form (BSF) 90-13 parasites were cultured at 37°C with 5% CO_2_ in modified HMI-9 culture medium with methylcellulose (MC-HMI-9), 10% heat-inactivated fetal bovine serum (FBS), 2.5 µg/mL G418, and 5 µg/mL hygromycin B. Slender BSF parasites were maintained below 7×10^5^ cells/mL to prevent stumpy BSF formation. Procyclic form (PCF) 90-13 parasites were cultured at 27°C in SDM80 culture medium supplemented with 10% heat-inactivated FBS, 50 mM N-acetyl-D-glucosamine, 15 µg/mL G418, and 50 µg/mL hygromycin B. To detect RDK1, we endogenously tagged a single chromosomal locus of RDK1 (TriTrypDB gene ID: Tb927.11.14070) with a C-terminal 10xTy epitope tag using the long PCR transfection method using a modified pPOTv4-10xTy plasmid containing a blasticidin S resistance (*BsdR*) cassette as the template DNA (51). Slender BSF parasites were transfected as previously described, and positive transfectants were selected in culture by addition of 5 µg/mL blasticidin S. A similar method was used to endogenously tag RDK1 with a C-terminal V5-miniTurbo biotin ligase (RDK1-miniTurbo) where the 10xTy epitope DNA sequence in the pPOTv4-10xTy plasmid was replaced with the DNA sequence encoding V5-tagged miniTurbo. To generate RDK1 RNAi parasites, the p2T7-177 RDK1 RNAi plasmid that was previously generated (30) was linearized with NotI, transfected into BSF RDK1-10xTy parasites, and positive transfectants were selected with 2.5 µg/mL phleomycin.

### *In vitro* differentiation of bloodstream form parasites to the procyclic form

Slender BSF parasites were resuspended in fresh MC-HMI-9 at an initial cell density of 2×10^5^ cells/mL and cultured at 37°C with 5% CO_2_. After 24 hours, the BSF culture was replenished with 10 mM D-glucose, and the BSF culture were grown for an additional 24 hours without dilution to complete stumpy BSF formation. Stumpy BSF parasites were resuspended in cool (∼21°C) modified DTM supplemented 3 mM citrate/3 mM *cis*-aconitate (CCA) at a density of 2×10^6^ cells/mL and were cultured at 27°C with 5% CO_2_ for 24 hours. After 24 hours in DTM with CCA, parasites were collected and transferred to fresh DTM without CCA added for another 24 hours. Finally, the differentiated parasites were collected and resuspended in fresh SDM80 medium at a density of 2×10^6^ cells/mL and allowed to outgrow as healthy PCF.

### RNA isolation and quantitative PCR analysis

To isolate RNA, we collected 1×10^8^ cells and lysed in 1 mL of TRIzol™ Reagent (Invitrogen, Cat. No: 15596-018). To serve as an internal spike-in control, we added 2 µL of diluted luciferase RNA (1 ng/µL) to the 1-mL TRIzol-cell lysate. Total RNA was purified by phenol-chloroform extraction and ethanol precipitation. Approximately 10 µg of RNA was rigorously treated with 6 U of TURBO DNase™ (Invitrogen, Cat. No: AM1907) for 90 minutes at 37°C to remove DNA contamination and re-purified by phenol-chloroform extraction and ethanol precipitation. DNase-treated RNA was converted to cDNA with the iScript™ cDNA Synthesis Kit (Bio-Rad, Cat. No: 1708891) using random hexamer primers. Quantitative PCR (qPCR) amplification and detection was performed on a CFX Real Time System thermocycler (Bio-Rad), and data were analyzed using the CFX Maestro software (Bio-Rad). The nucleotide sequences of RDK1 qPCR primers are included in Text S1. All qPCR experiments comprised of three, independent biological replicates of which each had three technical replicates. Values reported in this study are the arithmetic means of the measured values of the biological replicates with their corresponding standard error of the means.

### SDS-PAGE and Western blotting

To prepare whole cell lysates, we collected 5×10^7^ parasites, washed once in 1 mL of phosphate-buffered saline (PBS), and lysed in 300 µL of cell lysis buffer (20 mM Bis-Tris pH 7.5, 50 mM NaCl, 10 mM EDTA pH 8.0, 1.5% v/v Triton X-100, 0.2% w/v SDS, cOmplete™ EDTA-free Protease Inhibitor Cocktail [Roche/Millipore Sigma, Cat. No: 11873580001], and 0.1% w/v glycerol). Cell lysates were incubated at room temperature for 5 minutes before the addition of 150 µL of Bolt™ LDS Sample Buffer (Invitrogen, Cat. No: B0007) and 50 µL of 10X Bolt™ Sample Reducing Agent (Invitrogen, Cat. No: B0009). The completed samples were thoroughly vortexed to mix and heated at 90°C for 5 minutes. For SDS-PAGE analysis, 10 – 20 µL of parasite lysate (1 – 2×10^6^ cell equivalents) were loaded on a 4-12% Bolt™ Bis-Tris Plus Mini Protein gel (Invitrogen, Cat. No: NW04122BOX or NW04127BOX) and electrophoresed in 1X Bolt™ MOPS SDS Running Buffer (Invitrogen, Cat. No: B0001) at a constant 180 V for 20 minutes per manufacturer instructions. Proteins were transferred to a nitrocellulose membrane, blocked with EveryBlot Blocking Buffer (Bio-Rad, Cat. No: 12010020) and probed with primary antibodies or probes diluted in 1X Tris-buffered saline containing 0.25% Tween-20. For detecting biotinylated protein in RDK1-miniTurbo cell lysates, we probed with 1 µg/mL streptavidin-conjugated horseradish peroxidase (HRP) (Thermo Fisher, Cat No: S911) as a primary probe, washed three times with TBST, and detected chemiluminescence by SuperSignal™ West Femto Maximum Sensitivity Substrate (Thermo Fisher, Cat. No: 34096) using the ChemiDoc Go Imaging System (Bio-Rad, Cat. No: 12018025). For all other proteins, we used Goat Anti-Mouse StarBright™ Blue 520 and Goat Anti-Rabbit StarBright™ Blue 700 (Bio-Rad, Cat. No: 12005866 and 12004161) as secondary antibodies, and protein signals were detected using the ChemiDoc Go Imaging System. The full list of primary antibodies, secondary antibodies, their corresponding sources, and dilutions are listed in Text S1.

### Chemical modification of streptavidin beads and proximity-dependent protein biotinylation and identification

Dynabeads MyOne Streptavidin C1 magnetic beads (ThermoFisher: Cat. No: 65001) were chemically modified to confer trypsin resistance as previously described (52). BSF *T. brucei* parasites expressing RDK1 fused with a C-terminal V5-miniTurbo from an endogenous locus were cultured in fresh MC-HMI-9 supplemented with 50 µM biotin for 12 hours. As a negative control, parental BSF 90-13 parasites were treated under identical experimental conditions. Approximately 5×10^8^ parasites were collected, lysed in 5 mL of ice-cold lysis buffer (50 mM Tris-HCl pH 7.5, 500 mM NaCl, 0.4% w/v SDS, 5 mM EDTA pH 8.0, 1 mM DTT, 1X Protease Inhibitor Cocktail, 2% v/v Triton X-100), incubated on ice for 30 minutes, and the lysate was clarified by centrifuging at 16,000xg at 4°C for 30 minutes. Clarified lysate was incubated with 150 µL of chemically modified MyOne Streptavidin C1 Dynabeads for 2 hours at 4°C with constant rotation. After incubation, beads were washed twice with lysis buffer, twice with wash buffer 1 (2% w/v SDS), twice with wash buffer 2 (50 mM Tris-HCl pH 7.5, 0.5% v/v NP-40, 0.1% w/v sodium deoxycholate, 250 mM LiCl, and 1 mM EDTA), and twice with wash buffer 3 (50 mM Tris-HCl pH 7.5, 0.1% v/v NP-40, 500 mM NaCl). Samples were resuspended in 150 µL in wash buffer 3 and stored at -80°C until samples were sent to Cornell University’s Biotechnology Resource Center Proteomics and Metabolomics Facility for mass spectrometry identification of biotinylated proteins enriched in RDK1-miniTurbo parasites compared to parental 90-13 parasites.

### Quantitative global proteomics and phosphopeptide enrichment

For each sample, we collected 5×10^8^ parasites and washed once with 1 mL of ice-cold PBS containing 5 mM EDTA. We subsequently resuspended the parasites in 600 µL of ice-cold hypotonic lysis buffer (4X Halt™ Protease and Phosphatase Inhibitor Cocktail [ThermoFisher, Cat. No: 78441] and 1 mM EDTA pH 8.0), briefly vortexed, and let incubate at room temperature for 5 minutes. Following hypotonic lysis, we added 400 µL of solubilization buffer (10% w/v SDS, 500 mM DTT). Solubilized samples were stored at -80°C until analysis. Four independent biological replicates of BSF uninduced and induced RDK1 RNAi samples were generated and sent to Cornell University’s Biotechnology Resource Center Proteomics and Metabolomics Facility for mass spectrometry identification and parallel quantitative global proteomics and phosphoproteomics analyses. For global proteomics, 20 µg of protein from each sample was digested with trypsin. For quantitative phosphoproteomics, a total of 1,480 µg of protein was digested with trypsin, and phosphopeptides were enriched and analyzed by data-independent acquisition with label-free quantitation (DIA-LFQ).

### Mitochondrial membrane potential measurement

To measure mitochondrial membrane potential, 1×10^7^ of BSF uninduced control, BSF induced RDK1 RNAi, and wildtype PCF parasites were suspended in serum-free culture medium with 50 nM tetramethylrhodamine ethyl ester (TMRE) for 30 minutes at 37°C (for BSF) or 27°C (for PCF). As a control for depolarization of mitochondrial membrane potential, parallel samples of wildtype PCF parasites pre-treated with 10 µM carbonyl cyanide 4-(trifluoromethoxy)phenylhydrazone (FCCP) for 10 minutes prior to TMRE staining. Parasites were washed twice with cold 1X PBSG (supplemented with 5.5 mM glucose) immediately analyzed by flow cytometry (FACS Celesta). 10,000 fluorescent events were collected, and the geometric mean fluorescence intensity (gMFI) was calculated for each sample using the FlowJo software.

### Quantitation of mitochondrial superoxide levels

To measure mitochondrial superoxide levels, 1×10^7^ of BSF uninduced control, BSF induced RDK1 RNAi, and wildtype PCF parasites were suspended in serum-free culture medium with 5 µM MitoSOX™ Red Mitochondrial Superoxide Indicator (ThermoFisher, Cat. No: M36007) for 30 minutes at 37°C (for BSF) or 27°C (for PCF). As a control for the induction of mitochondrial superoxide, parallel samples of wildtype PCF parasites pre-treated with 100 µM paraquat for 30 minutes prior to MitoSOX™ staining. Parasites were washed twice with cold 1X PBSG (supplemented with 5.5 mM glucose) immediately analyzed by flow cytometry (FACS Celesta). 10,000 fluorescent events were collected, and the geometric mean fluorescence intensity (gMFI) was calculated for each sample using the FlowJo software.

### *In vitro* kinase and adenyl cyclase assays

We collected 5×10^8^ BSF parental 90-13 parasites or parasites expressing RDK1 with a C-terminal 10xTy epitope tag (RDK1-10xTy), lysed in 1 mL of cold lysis buffer (10 mM Bis-Tris pH 7.5, 50 mM NaCl, 4X Halt™ Protease and Phosphatase Inhibitor Cocktail [ThermoFisher, Cat. No: 78441], 5 mM EDTA pH 8.0, and 1% v/v Triton X-100), and incubated the lysate on ice for 30 minutes. We clarified the lysate by centrifuging at 18,000xg at 4°C for 15 minutes. The clarified lysate was then incubated with 50 µL of Protein A-Sepharose FastFlow beads (Millipore Sigma, Cat. No: P9424-1ML) conjugated to anti-Ty antibodies for 2 hours while rotating at 4°C.

Beads were washed twice with cold wash buffer (10 mM Bis-Tris pH 7.5, 50 mM NaCl, 1X Halt™ Protease and Phosphatase Inhibitor Cocktail, 5 mM EDTA pH 8.0, and 0.1 % v/v Triton X-100). For kinase assays, beads with immunoprecipitated proteins were resuspended in kinase buffer (50 mM Tris-HCl pH 7.5, 10 mM MgCl_2_, 1 mM DTT, and 0.1 mg/mL bovine serum albumin). ATP was added to a final concentration of 50 µM before kinase activity was measured using the ADP-Glo™ Kinase Assay (Promega, Cat. No: PAV6930) according to manufacturer’s protocol. For adenyl cyclase assays, beads with immunoprecipitated proteins were resuspended in 15 µL of induction buffer made in 1X PBS (0.5 mM 3-isobutyl-1-methylxanthine, 0.1 mM Ro-20-1724) and added 5 µL of 1X PBS supplemented with 400 µM ATP. Using cAMP-Glo™ Assay kit reagents (Promega, Cat. No: V1501), 20 µL of cAMP-Glo™ lysis buffer and incubated at room temperature for 15 minutes. 40 µL of cAMP Detection solution were added and the samples were agitated for 1 minute before incubation at room temperature for 20 minutes. 80 µL of Kinase-Glo Reagent supplemented with protein kinase A (PKA) was added and incubated at room temperature for 10 minutes. Each sample was split into three technical replicates. Buffer controls containing 0, 1, 10, or 100 nM cAMP were included, and cAMP levels were measured by a luminometer.

### Intracellular cAMP measurement

To measure intracellular cyclic AMP (cAMP) levels, ∼3.2×10^7^ parasites were washed once with cold 1X PBS, resuspended in 20 µL of induction buffer made with sterile water (0.5 mM 3-isobutyl-1-methylxanthine, 0.1 mM Ro-20-1724), and incubated at 30°C for 15 minutes to allow for complete hypotonic lysis. Using cAMP-Glo™ Assay kit reagents (Promega, Cat. No: V1501), 20 µL of cAMP-Glo™ lysis buffer and incubated at room temperature for 15 minutes. 40 µL of cAMP Detection solution were added and the samples were agitated for 1 minute before incubation at room temperature for 20 minutes. 80 µL of Kinase-Glo Reagent supplemented with protein kinase A (PKA) was added and incubated at room temperature for 10 minutes. Each sample was split into three technical replicates, and cAMP levels were measured by a luminometer.

## ACKNOWLEDGMENTS

We greatly acknowledge the team at the Biotechnology Research Center Proteomics Team at Cornell University for their assistance with proteomics and phosphoproteomics analyses. We thank Dr. James Bangs at the University at Buffalo for providing anti-VSG and anti-EP1 antibodies. We thank Dr. Christopher de Graffenried from Brown University for providing anti-Ty1 antibody.

---

**Text S1.**
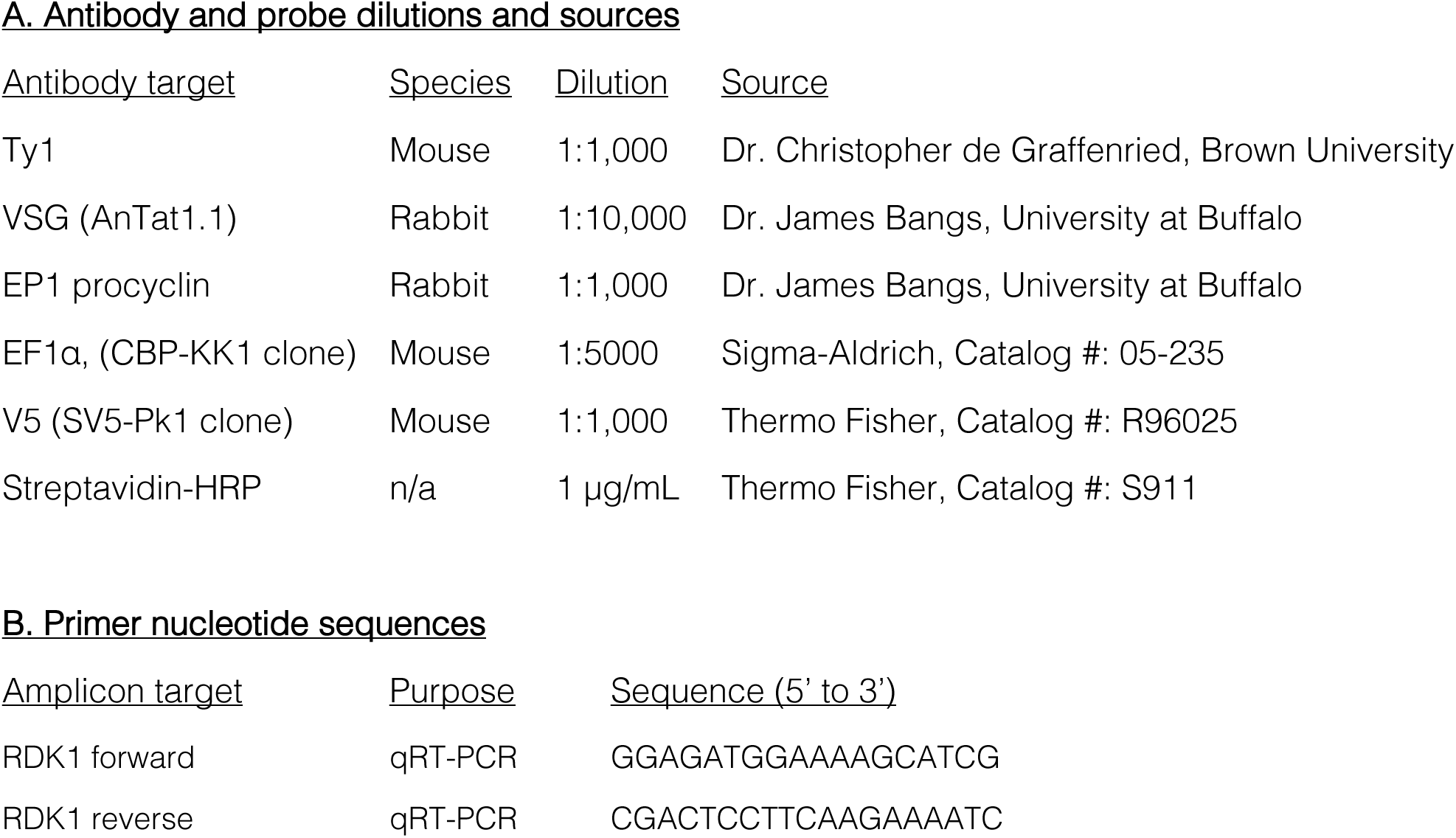
List of primers and antibodies.

**Table S1.** Complete list of RDK1-proximal proteins in BSF parasites

**Table S2.** Complete list of upregulated proteins in RDK1 RNAi parasites

**Table S3.** Complete list of differentially abundant phosphosites in RDK1 RNAi parasites

## REFERENCES

1. Lundkvist GB, Kristensson K, Bentivoglio M. 2004. Why trypanosomes cause sleeping sickness. Physiology 19:198–206.

2. Desquesnes M, Gonzatti M, Sazmand A, Thévenon S, Bossard G, Boulangé A, Gimonneau G, Truc P, Herder S, Ravel S, Sereno D, Jamonneau V, Jittapalapong S, Jacquiet P, Solano P, Berthier D. 2022. A review on the diagnosis of animal trypanosomoses. Parasites Vectors 15:64.

3. Szöőr B, Silvester E, Matthews KR. 2020. A leap into the unknown – early events in African trypanosome transmission. Trends Parasitol 36:266–278.

4. Mogk S, Meiwes A, Shtopel S, Schraermeyer U, Lazarus M, Kubata B, Wolburg H, Duszenko M. 2014. Cyclical appearance of African trypanosomes in the cerebrospinal fluid: New insights in how trypanosomes enter the CNS. PLoS ONE 9:e91372.

5. Trindade S, Rijo-Ferreira F, Carvalho T, Pinto-Neves D, Guegan F, Aresta-Branco F, Bento F, Young SA, Pinto A, Van Den Abbeele J, Ribeiro RM, Dias S, Smith TK, Figueiredo LM. 2016. *Trypanosoma brucei* parasites occupy and functionally adapt to the adipose tissue in mice. Cell Host Microbe 19:837–848.

6. Trindade S, Niz MD, Costa-Sequeira M, Bizarra-Rebelo T, Bento F, Dejung M, Narciso MV, López-Escobar L, Ferreira J, Butter F, Bringaud F, Gjini E, Figueiredo LM. 2022. Slow growing behavior in African trypanosomes during adipose tissue colonization. Nat Commun 13:7548.

7. Mabille D, Dirkx L, Thys S, Vermeersch M, Montenye D, Govaerts M, Hendrickx S, Takac P, Weyenbergh JV, Pintelon I, Delputte P, Maes L, Pérez-Morga D, Timmermans J-P, Caljon G. 2022. Impact of pulmonary African trypanosomes on the immunology and function of the lung. Nat Commun 13:7083.

8. Rico E, Rojas F, Mony BM, Szoor B, MacGregor P, Matthews KR. 2013. Bloodstream form pre-adaptation to the tsetse fly in *Trypanosoma brucei*. Front Cell Infect Microbiol 3:78.

9. Matthews KR. 2005. The developmental cell biology of *Trypanosoma brucei*. J Cell Sci 118:283–290.

10. Silvester E, McWilliam KR, Matthews KR. 2017. The cytological events and molecular control of life cycle development of *Trypanosoma brucei* in the mammalian bloodstream. Pathogens 6:29.

11. Qiu Y, Milanes JE, Jones JA, Noorai RE, Shankar V, Morris JC. 2018. Glucose signaling is important for nutrient adaptation during differentiation of pleomorphic African trypanosomes. mSphere 3:e00366–18.

12. Rojas F, Silvester E, Young J, Milne R, Tettey M, Houston DR, Walkinshaw MD, Pérez-Pi I, Auer M, Denton H, Smith TK, Thompson J, Matthews KR. 2019. Oligopeptide signaling through TbGPR89 drives trypanosome quorum sensing. Cell 176:306–317.e16.

13. Szöőr B, Ruberto I, Burchmore R, Matthews KR. 2010. A novel phosphatase cascade regulates differentiation in *Trypanosoma brucei* via a glycosomal signaling pathway. Genes Dev 24:1306–1316.

14. Mony BM, MacGregor P, Ivens A, Rojas F, Cowton A, Young J, Horn D, Matthews K. 2013. Genome-wide dissection of the quorum sensing signalling pathway in *Trypanosoma brucei*. Nature 505:681–685.

15. Szöőr B, Dyer NA, Ruberto I, Acosta-Serrano A, Matthews KR. 2013. Independent pathways can transduce the life-cycle differentiation signal in *Trypanosoma brucei*. PLoS Pathog 9:e1003689.

16. McDonald L, Cayla M, Ivens A, Mony BM, MacGregor P, Silvester E, McWilliam K, Matthews KR. 2018. Non-linear hierarchy of the quorum sensing signalling pathway in bloodstream form African trypanosomes. PLoS Pathog 14:e1007145.

17. Jones NG, Thomas EB, Brown E, Dickens NJ, Hammarton TC, Mottram JC. 2014. Regulators of *Trypanosoma brucei* cell cycle progression and differentiation identified using a kinome-wide RNAi screen. Plos Pathog 10:e1003886.

18. Benz C, Urbaniak MD. 2019. Organising the cell cycle in the absence of transcriptional control: Dynamic phosphorylation co-ordinates the *Trypanosoma brucei* cell cycle post-transcriptionally. PLoS Pathog 15:e1008129.

19. Domingo -Sananes MR, Szöőr B, Ferguson MAJ, Urbaniak MD, Matthews KR. 2015. Molecular control of irreversible bistability during trypanosome developmental commitment. J Cell Biol 211:455–468.

20. Ooi CP, Benz C, Urbaniak MD. 2020. Phosphoproteomic analysis of mammalian infective *Trypanosoma brucei* subjected to heat shock suggests atypical mechanisms for thermotolerance. J Proteom 219:103735.

21. Toh JY, Nkouawa A, Sánchez SR, Shi H, Kolev NG, Tschudi C. 2021. Identification of positive and negative regulators in the stepwise developmental progression towards infectivity in *Trypanosoma brucei*. Sci Rep-uk 11:5755.

22. Pfister DD, Burkard G, Morand S, Renggli CK, Roditi I, Vassella E. 2006. A mitogen-activated protein kinase controls differentiation of bloodstream forms of *Trypanosoma brucei*. Eukaryot Cell 5:1126–1135.

23. Cayla M, McDonald L, MacGregor P, Matthews K. 2020. An atypical DYRK kinase connects quorum-sensing with posttranscriptional gene regulation in *Trypanosoma brucei*. eLife 9:e51620.

24. Jensen BC, Vaney P, Flaspohler J, Coppens I, Parsons M. 2021. Unusual features and localization of the membrane kinome of *Trypanosoma brucei*. PLoS ONE 16:e0258814.

25. Caveney NA, Tsutsumi N, Garcia KC. 2023. Structural insight into guanylyl cyclase receptor hijacking of the kinase–Hsp90 regulatory mechanism. eLife 12:RP86784.

26. Potter LR. 2024. Phosphorylation-dependent regulation of guanylyl cyclase (GC)-A and other membrane GC receptors. Endocr Rev 45:755–771.

27. Bhandari R, Srinivasan N, Mahaboobi, Ghanekar Y, Suguna K, Visweswariah SS. 2001. Functional inactivation of the human guanylyl cyclase C receptor: Modeling and mutation of the protein kinase-like domain. Biochemistry 40:9196–9206.

28. Jaleel M, Saha S, Shenoy AR, Visweswariah SS. 2006. The kinase homology domain of receptor guanylyl cyclase C: ATP binding and identification of an adenine nucleotide sensitive site. Biochemistry 45:1888–1898.

29. Bose A, Visweswariah SS. 2022. The pseudokinase domain in receptor guanylyl cyclases. Methods Enzym 667:535–574.

30. Smith JT, Tylec BL, Naguleswaran A, Roditi I, Read LK. 2023. Developmental dynamics of mitochondrial mRNA abundance and editing reveal roles for temperature and the differentiation-repressive kinase RDK1 in cytochrome oxidase subunit II mRNA editing. mBio.

31. Jensen BC, Ramasamy G, Vasconcelos EJR, Ingolia NT, Myler PJ, Parsons M. 2014. Extensive stage-regulation of translation revealed by ribosome profiling of *Trypanosoma brucei*. BMC Genom 15:911.

32. Naguleswaran A, Doiron N, Roditi I. 2018. RNA-Seq analysis validates the use of culture-derived *Trypanosoma brucei* and provides new markers for mammalian and insect life-cycle stages. BMC Genom 19:227.

33. Dejung M, Subota I, Bucerius F, Dindar G, Freiwald A, Engstler M, Boshart M, Butter F, Janzen CJ. 2016. Quantitative proteomics uncovers novel factors involved in developmental differentiation of *Trypanosoma brucei*. PLoS Pathog 12:e1005439.

34. Dewar CE, Oeljeklaus S, Mani J, Mühlhäuser WWD, Känel C von, Zimmermann J, Ochsenreiter T, Warscheid B, Schneider A. 2022. Mistargeting of aggregation prone mitochondrial proteins activates a nucleus-mediated posttranscriptional quality control pathway in trypanosomes. Nat Commun 13:3084.

35. Singha UK, Hamilton V, Chaudhuri M. 2015. Tim62, a novel mitochondrial protein in *Trypanosoma brucei*, is essential for assembly and stability of the TbTim17 protein complex. J Biol Chem 290:23226–23239.

36. Fernandez-Cortes F, Serafim TD, Wilkes JM, Jones NG, Ritchie R, McCulloch R, Mottram JC. 2017. RNAi screening identifies *Trypanosoma brucei* stress response protein kinases required for survival in the mouse. Sci Rep 7:6156.

37. Hammarton TC. 2007. Cell cycle regulation in *Trypanosoma brucei*. Mol Biochem Parasitol 153:1–8.

38. Hu H, Majneri P, Li D, Kurasawa Y, An T, Dong G, Li Z. 2017. Functional analyses of the CIF1–CIF2 complex in trypanosomes identify the structural motifs required for cytokinesis. J Cell Sci 130:4108–4119.

39. Saldivia M, Wollman AJM, Carnielli JBT, Jones NG, Leake MC, Bower-Lepts C, Rao SPS, Mottram JC. 2021. A CLK1-KKT2 signaling pathway regulating kinetochore assembly in *Trypanosoma brucei*. mBio 12:10.1128/mbio.00687-21.

40. Cayla M, Nievas YR, Matthews KR, Mottram JC. 2022. Distinguishing functions of trypanosomatid protein kinases. Trends Parasitol 38:950–961.

41. Ballmer D, Akiyoshi B. 2024. Dynamic localization of the chromosomal passenger complex is controlled by the orphan kinesins KIN-A and KIN-B in the kinetoplastid parasite *Trypanosoma brucei*. eLife 10.7554/elife.93522.2.

42. Doleželová E, Kunzová M, Dejung M, Levin M, Panicucci B, Regnault C, Janzen CJ, Barrett MP, Butter F, Zíková A. 2020. Cell-based and multi-omics profiling reveals dynamic metabolic repurposing of mitochondria to drive developmental progression of *Trypanosoma brucei*. Plos Biol 18:e3000741.

43. Docampo R, Vercesi AE. 2022. Mitochondrial Ca^2+^ and reactive oxygen species in trypanosomatids. Antioxid Redox Signal 36:969–983.

44. Badjatia N, Ambrósio DL, Lee JH, Günzl A. 2013. Trypanosome cdc2-related kinase 9 controls spliced leader RNA *cap4* methylation and phosphorylation of RNA polymerase II subunit RPB1. Mol Cell Biol 33:1965–1975.

45. Hammarton TC, Kramer S, Tetley L, Boshart M, Mottram JC. 2007. *Trypanosoma brucei* polo-like kinase is essential for basal body duplication, kDNA segregation and cytokinesis. Mol Microbiol 65:1229–1248.

46. Carlson J, Ahmed M, Hunter R, Hoque SF, Benoit JB, Chiurillo MA, Lander N. 2025. TcCARP3 modulates compartmentalized cAMP signals involved in osmoregulation, infection of mammalian cells, and colonization of the triatomine vector in the human pathogen *Trypanosoma cruzi*. mBio 16:e00994–25.

47. Panigrahi AK, Ogata Y, Zíková A, Anupama A, Dalley RA, Acestor N, Myler PJ, Stuart KD. 2009. A comprehensive analysis of *Trypanosoma brucei* mitochondrial proteome. Proteomics 9:434–450.

48. Taleva G, Husová M, Panicucci B, Hierro-Yap C, Pineda E, Biran M, Moos M, Šimek P, Butter F, Bringaud F, Zíková A. 2023. Mitochondrion of the *Trypanosoma brucei* long slender bloodstream form is capable of ATP production by substrate-level phosphorylation. PLOS Pathog 19:e1011699.

49. Dai Y, Schlanger S, Haque MM, Misra S, Stuehr DJ. 2019. Heat shock protein 90 regulates soluble guanylyl cyclase maturation by a dual mechanism. J Biol Chem 294:12880–12891.

50. Engstler M, Boshart M. 2004. Cold shock and regulation of surface protein trafficking convey sensitization to inducers of stage differentiation in *Trypanosoma brucei*. Genes Dev 18:2798–2811.

51. Dean S, Sunter J, Wheeler RJ, Hodkinson I, Gluenz E, Gull K. 2015. A toolkit enabling efficient, scalable and reproducible gene tagging in trypanosomatids. Open Biol 5:140197.

52. Rafiee M, Sigismondo G, Kalxdorf M, Förster L, Brügger B, Béthune J, Krijgsveld J. 2020. Protease-resistant streptavidin for interaction proteomics. Mol Syst Biol 16:MSB199370.

